# Anti-malaria RTS,S/AS01 vaccine generates a limited pool of memory B cells expressing specificities associated with protection

**DOI:** 10.64898/2025.12.22.696031

**Authors:** Jason Netland, Christopher D. Thouvenel, Scott Gregory, William C. Adams, Erik Jongert, Natalie Brunette, Neil P. King, Neville K. Kisalu, C. Richter King, David J. Rawlings, Marion Pepper

**Author notes:** Authors contributed equally to this project (Jason Netland, Christopher D. Thouvenel, David J. Rawlings, Marion Pepper).

## Abstract

RTS,S/AS01, an adjuvanted subunit malaria vaccine, induces protective but short-lived anti- *Plasmodium falciparum* circumsporozoite protein (CSP) antibody titers. To better understand the lack of sustained protection post-vaccination, we analyzed CSP-specific B cells over time in malaria-naïve individuals following immunization with RTS,S/AS01. Longitudinal analyses of the cellular responses revealed a shift in the specificity of CSP-specific B cells over time. Early post-vaccine responses were dominated by plamsablasts and class-switched memory B cells (MBCs) specific for the NANP-repeat region of CSP, epitopes associated with protective antibodies. However, the frequency of NANP-repeat-specific MBCs declines, while longer-term memory specific for the C-terminus of CSP persists. Despite being class-switched and affinity matured, C-term monoclonal antibodies derived from these MBCs failed to protect mice against a transgenic parasite challenge. Taken together, these findings suggest that RTS,S/AS01 induces a transient, protective NANP repeat-specific B cell response that is subsequently replaced by memory B cells with non-protective reactivities, potentially explaining its limited long-term efficacy and limited response to subsequent challenges or boosters.

## Introduction

Malaria remains a major global threat with an estimated 282 million cases and 610,000 deaths in 2024 alone^1^. The currently available malaria vaccines, RTS,S/AS01 and R21/Matrix-M, are subunit vaccines that contain components of *Plasmodium falciparum* circumsporozoite protein (CSP) antigen, the major protein on the surface of the infective stage of the malaria parasite. Both vaccines contain a portion of CSP displayed on a HepB virus-like particle. Antibodies against CSP have been shown to be highly protective in humans, and passive transfer of an anti-CSP junctional antibody can protect against human infection for months^2–4^. Despite this potential for humoral protection, CSP-based vaccine efficacy is limited and protection begins to wane within months^5–8^, correlated with a loss of protective serum antibodies^9^

In an effort to better understand the RTS,S-induced B cell response over time, we examined CSP-specific B cells in the blood of malaria-naïve volunteers over the course of vaccination and controlled human malarial infection (CHMI) trials. RTS,S contains an N-terminally truncated PfCSP protein consisting of a shortened NANP-repeat and the entire C-terminus (C-term) of CSP^10^. Previous work by numerous groups has sought to determine the relative contributions of serum antibody reponses targeting these distinct regions to the protection elicted following vaccination^9,11–16^. While it is well appreciated that repeat-region specific antibodies can be protective^17^, it has also been suggested that this region may serve as a “decoy” to prevent development of higher affinity responses to other regions of the protein^18,19^. Work in mouse models indicates that the highly repetitive nature of this epitope allows it to dominate the response and that shortening the repeat region increases reponses to other regions of CSP^20^. However, only a few C-term specific antibodies have been isolated from human subjects, and they either fail to protect or do so only modestly^21–23^. To date, a longitudinal analysis of the B cell responses to these regions in humans following vacination with RTS,S has not been carried out. To this end, we used B cell tetramers targeting these functionally distinct regions of CSP to isolate and analyze single B cells and their B cell receptors in humans following vaccination in order to determine their respective contributions to anti-CSP B cell responses.

Our studies focused on individuals from non-endemic regions in an effort to visualize the kinetics of the B cell response initiated in naïve individuals. In 2013, GlaxoSmithKline (GSK), PATH, and Walter Reed Army Instutute of Research (WRAIR) conducted the MALARIA-071 (MAL071) trial immunizing malaria-naïve participants with RTS,S/AS01_B_ on a 0, 1, 7-month schedule. This trial included subjects recieving a delayed “fractional”, 1/5^th^ dose at 7 months designed to enhance efficacy relative to a standard immunization schedule at 0, 1 and 2 months at full doses^10,24^. Following the primary phase of the MAL071 study, an additional, malaria parasite challenge 8 months after first CHMI provided a pilot assessment of whether protection was maintained and whether the fractional dose might convey higher levels of sterile immunity. Following our initial assessment of samples from MAL071, we also analyzed samples from subjects in a follow-up study, MALARIA-092 (MAL092)^25^ that utilized a vaccination and challenge schedule similar to the primary phase of MAL071. We employed a combination of antigen-specific flow-based B cell analysis and sorting, B cell receptor (BCR) sequencing, monoclonal antibody (mAb) generation and functional antibody characterization to examine the quantity, quality and specificity of CSP-reactive B cells generated by RTS,S/AS01_B_ vaccination over time.

Our study reveals that CSP-specific B cell responses immediately following RTS,S/AS01 vaccination are dominated by class-switched cells that recognize the NANP-repeat region of CSP, specificities known to be protective against infection. However these NANP-repeat-specific cells appear to be short-lived, and the long-lived memory pool is instead dominated by class-switched C-term-specific memory B cells. Despite these C-term-specific cells expressing affinity-matured BCRs, their antibodies do not appear to be protective in an intravenous (IV) challenge model, as has been found for other mAbs reactive to the C-term^21,22^. Taken together, our findings may help to explain the relatively short period of protection and limited protective response to booster doses associated with RTS,S and suggest that alternative approaches are likely to be required to generate long-lived, NANP-repeat-specific, protective B cell responses.

## Results

### Isolation and sequencing of CSP-specific B cells from vaccinated individuals

To determine if it was feasible to identify, isolate and analyze CSP-specific B cells responding to vaccination, we obtained peripheral blood mononuclear cells (PBMCs) from two protected individuals enrolled in the MAL071 RTS,S/AS01 vaccine trial. PBMCs were collected prior to vaccination (Timepoint A) and 7 days post-third vaccination (Timepoint B^7^) in accordance with the standard RTS,S/AS01 vaccination schedule (Fig. 1a). Samples were first labeled with a decoy tetramer followed by addition of PE-conjugated CSP tetramers to rigorously identify B cells that bind CSP^26^. Magnetic bead enrichment of PE-labeled B cells facilitated identification of rare CSP-specific B cells. Enriched cells were subsequently stained with a panel of antibodies against B cell surface proteins to identify distinct populations of B cells within single-cell index FACS-sorted CSP-specific B cells. CSP-specific B cells were phenotypically characterized using an antibody panel that included markers designed to characterize memory B cell populations (CD21/CD27), short-lived plasmablasts (CD38,CD27) and specific BCR isotypes (IgG/IgM/IgD/IgA) (Figs. 1b and Extended Data Fig. 1a). This approach allowed us to identify a small number of naïve CD21^+^CD27^-^ CSP-specific B cells in individuals pre-immunization (Fig. 1b). As hypothesized, the number of CSP-specific B cells increased substantially post-third immunization (5-7 fold compared to naïve cell numbers) (Fig. 1c). Expanded CSP-specific B cells expressed cell surface proteins associated with immune activation and most expressed markers consistent with differentiation into specific fates including memory B cells (CD21+CD27+ classical memory, CD21^+^CD27^-^ activated memory and CD21^-^CD27^-^ atypical memory), and short-lived plasmablasts (CD27^+^CD38^+^) (Fig. 1b).

**Fig. 1:**
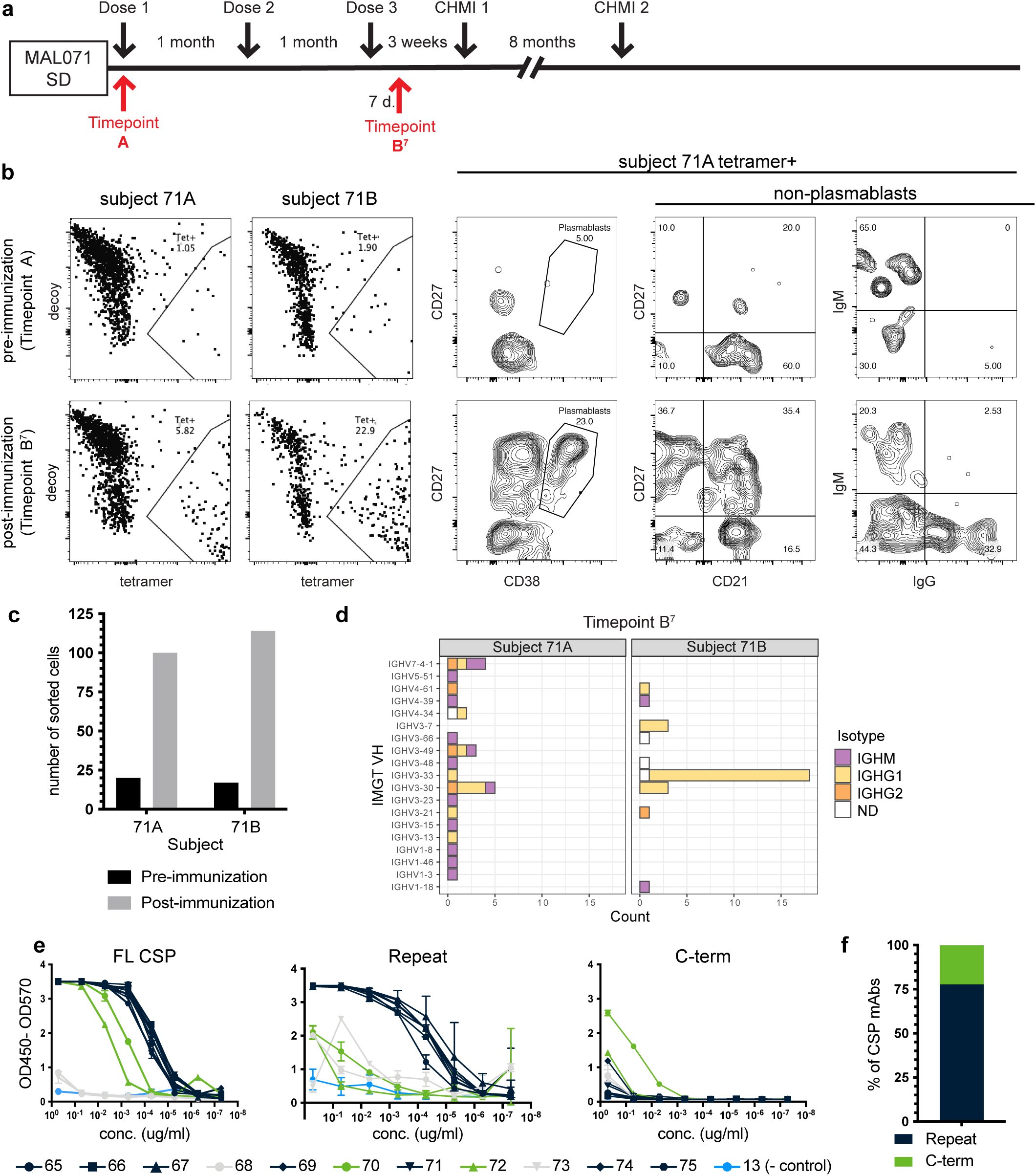
Detection of CSP- specific B cells in RTS,S vaccinated individuals. **a**, Timeline of vaccination/challenge regime for MAL-071 SD trial arm and PBMC sampling. PBMC samples analyzed were prior to vaccine dose 1 (Timepoint A) and 7 days post-3rd dose (Timepoint B7). **b,** Representative flow plots of sorted CSP-specific B cells, tetramer specific cells were further analyzed by cell surface marker stain to identify distinct populations such as memory B cells (CD27, CD21), plasmablasts (CD27^+^CD38^hi^) and antibody isotypes (IgG,IgM). **c,** Number of CSP tetramer positive cells sorted pre- and post- vaccination. **d,** Heavy chain gene usage per subject of sorted and sequenced B cells 7 days post 3 vaccine dose. Isotype indicated was determined by sequencing. IMGT, the international ImMunoGeneTics information system. IGHV, Immunoglobulin (Ig) heavy variable region gene family. Isotype: IGHM, IgM heavy; IGHG1, IgG1 heavy; IGHG2, IgG2 heavy; ND, not determined. **e,** Binding of purified anti-CSP mAbs by ELISA to full length (FL) CSP, NANP repeat or C-term peptides. A SARS-CoV2 spike specific antibody (13) was used as a negative control. ELISA were conducted in triplicate and plots are representative of 3 independent experiments. **f,** Quantificiation of antibody specificity measured by ELISA. The proportion of mAbs targeting the NANP-repeat or C-terminal domain of PfCSP is shown.

We next sequenced and expressed BCRs from a series of single CSP-specific cells to assess epitope specificity and learn more about population clonality. Eighty-four single cells (42 from each of 2 different subjects) at timepoint B^7^ (7 days post-third immunizartion) (Fig. 1a) were randomly selected for BCR sequencing. BCR sequencing was performed on individual sorted cells such that paired heavy and light chain sequences could be captured (as described in the Materials and Methods). Productively rearranged sequences were obtained for 57 heavy chains and 63 light chains. Analysis of heavy and light chain gene usage, somatic hypermutation and CDR3 sequences demonstrated that all but two isolated CSP-specific B cells were unique clones that were derived from distinct VDJ rearrangements (Extended Data Table 1). Heavy chain (HC) gene usage revealed a broad set of genes in IgM expressing B cells including representation from variable heavy (VH) families 1, 3, 4, 5 and 7. BCRs utilizing an IgG HC exhibited a more restricted gene usage, only representing VH families 3, 4 and 7, with VH3-30 and VH3-33 being predominantly represented (Fig. 1d and Extended Data Fig. 1b). Notably, this IgG HC gene usage pattern was similar to that identified using alternative methods in recipients of either RTS,S or live attenuated sporozoite immunization^27,28^.

We next used the sequencing and phenotypic data from index sorted cells to select 11 BCRs to clone and recombinantly express to demonstrate CSP-specificity by ELISA. These BCRs were chosen to represent a diverse array of both HC families and different B cell subsets. Specifically, clones were selected based on a variety of isotypes (5 IgM, 6 IgG) , phenotypes (2 naïve, 3 short-lived plasmablasts, and 6 memory cells), and Ig-gene usage, with approximately 50% of total clones coming from each of the 2 subjects analyzed. Recombinant mAbs derived from BCR sequences from individual clones were tested by ELISA for specificity to the CSP antigen. To control for differences in avidity (IgM antibodies are multimers, while IgG antibodies are monomers) and differences in antibody detection (due to the use of different secondary antibodies required for each isotype), all recombinant mAbs were expressed with an IgG1 constant region regardless of the original isotype of the isolated B cell. Seven out of the 11 antibodies strongly bound CSP, while 2 antibodies bound slightly less well (Fig. 1e). We next determined which regions of the CSP each mAb could bind by conducting ELISAs using either a NANP-repeat region peptide (NANP6) or a C-terminus peptide (P16). All 7 of the antibodies that bound full-length CSP most strongly were specific to the NANP-repeat region, with 3 being derived from plasmablasts and 4 derived from memory B cells, while the 2 lower binders bound weakly to the C-term peptide (Fig. 1e,f). These data are consistent with numerous previous studies that have identified predominantly NANP-repeat-specific antibodies in individuals who have been exposed to CSP either through direct *Plasmodium* infection or RTS,S or sporozoite immunization^27,29–31^. Taken together, these findings validated our tetramer-based, single-cell approach for expanded studies of CSP-specific B cells derived from RTS,S-immunized individuals.

### Single-cell analysis of CSP-specific B cells from protected individuals in MAL071 trial

While malaria vaccines such as RTS,S have demonstrated consistent short-lived protection^32^, long-term protection and the ability to boost pre-exisiting responses have been more difficult to achieve for unknown reasons^32^. Based on our ability to isolate and characterize CSP-specific B cells following RTS,S vaccination at both the population and single-cell level, we next wanted to analyze the CSP-specific B cell response in the memory phase of the response. To address this question, we focused our efforts on analyzing B cell responses in the small number (n=3) of MAL071 trial participants that were protected following two CHMIs without receiving a vaccine booster^24^. Protected subjects were from the Delayed Fractional Dose arm of the study, having received 2 standard doses of the vaccine followed by a delayed, fractional third dose. Samples from 3 subjects across 4 timepoints, one at the time of second CHMI (timepoint C) and 3 subsequent timepoints at one month intervals post-second challenge (timepoints D, E and F), were analyzed (Fig. 2a). These timepoints were chosen to allow examination of the B cell memory pool present at the time of parasite challenge and any potential changes in this population in response to the challenge. CSP-specific B cells were isolated from PBMCs and analyzed as described above. Cells were first analyzed at the population level for differences in surface phenotype across the various timepoints. Approximately 70-90% of the CSP-specific cells expressed markers of memory, with a similar proportion of cells expressing class-switched IgG BCRs. Virtually no plasmablasts were dectected at these late time points, as expected. Examination of the frequencies of plasmablasts, memory B cells, and IgG^-^, IgM^-^, or IgA^-^expressing B cells revealed no major differences in individuals across the 4 timepoints (Fig. 2b and Extended Data Fig. 2a).

**Fig. 2:**
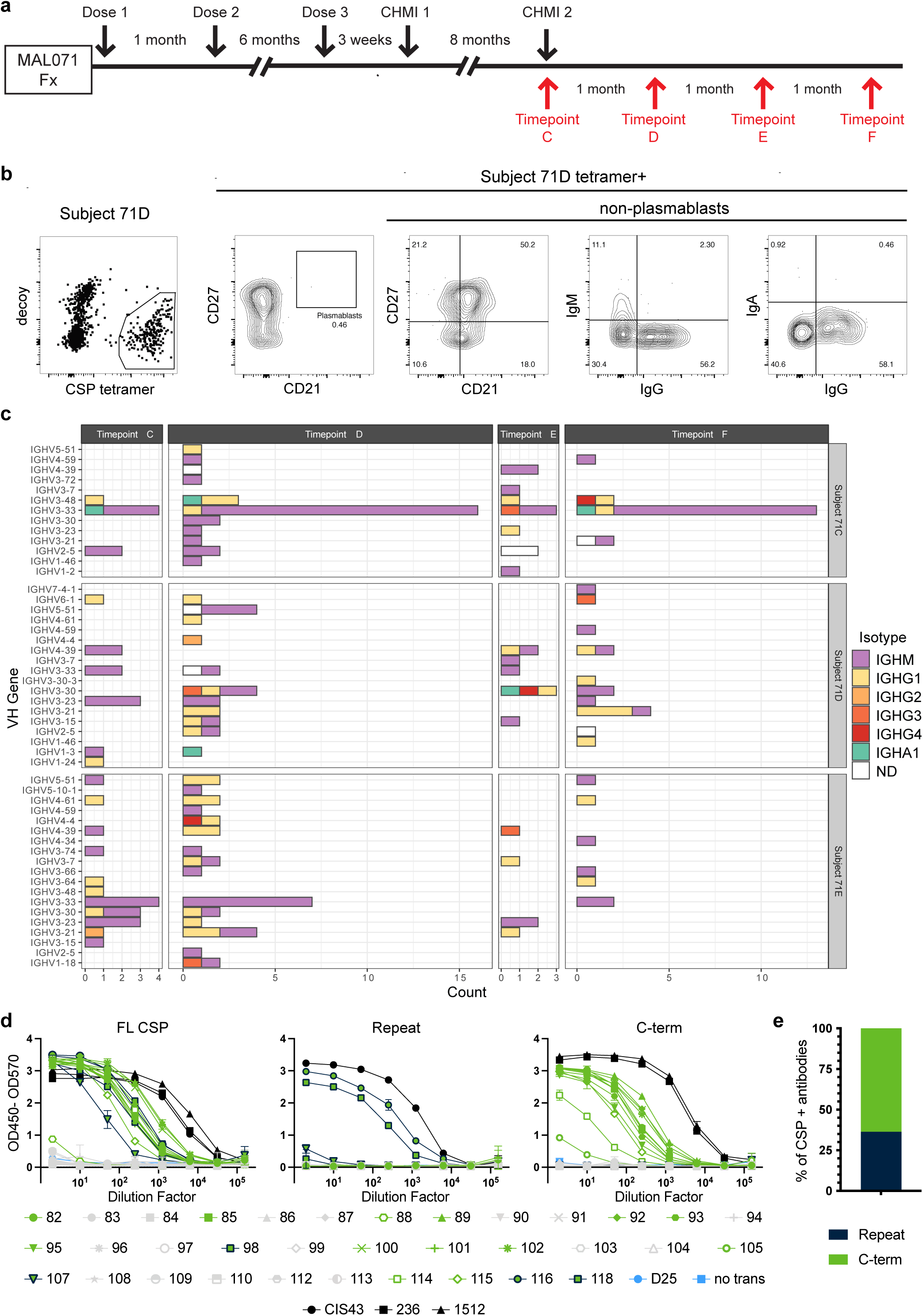
Memory B cell response in long-term protected indiviuals is dominated by C-term reactive cells. **a**, Timeline of vaccination/challenge regime for MAL-071 Fx trial arm and PBMC sampling. PBMC samples analyzed were from the day of 2nd CHMI (Timepoint C) and at approximately 1 month intervals out to 3 months (Timepoints D-F.) **b,** Example flow plots from sorts. Cell surface marker gates were set using non-antigen specific cells. Example shown is from day of 2nd CHMI (Timepoint C). **c,** BCR heavy chain gene usage of single CSP-specific B cells sorted from the 3 individuals who were protected from the 2nd CHMI. The isotype indicated was determined by sequencing. **d,** ELISAs of monoclonal antibody containing transfection supernatants showing binding to full length CSP, repeat and C-term peptides. A RSV-specific antibody (D25) and untransfected cells supernatant (no trans) were included as negative controls. A repeat specific antibody (CIS43) and C-term specific antibodies (236 and 1512) were included as positive controls. ELISA were conducted in triplicate and plots are representative of 3 independent experiments. **e,** Quantificiation of antibody specificity as detemined by ELISA.

We next single-cell sequenced CSP-specific B cells to determine BCR gene usage. Based on our initial data above and previous observations, we hypothesized that the majority of antigen-specific B cells at these timepoints would be comprised of memory phenotype, NANP-repeat-specific cells. A total of 266 CSP-specific cells were sorted across all three individuals (subject 71C, 71D, 71E) and timepoints. Heavy chain gene analysis revealed a potential bias for VH3-30, VH3-33, or VH3-23, previously shown to be associated with repeat/junction-specific antibodies^27,28,33^. However, this finding was primarily driven by a single individual and, surprisingly, was largely derived from IgM-expressing B cells in all individuals. Although IgG B cells expressing these VH genes were also present in the three subjects, overall IgG gene usage was much more diverse at all timepoints. (Fig. 2c). When light chains were included in the analysis, the diversity was even more evident as very few common heavy/light chain pairs were observed (Extended Data Fig. 2b), and with the exception of a single clone that was observed twice, all sequences were unique (Extended Data Table 2). Increased breadth in LC pairing was also evident in cells expressing the VH3-33 HC usage, previously shown to preferentially pair with Kappa 1-5 LC in NANP-repeat- specific antibodies^34,35^ ; an association not present in our dataset. Comparing the sequences we obtained to germline heavy chain sequences revealed that almost all the IgG and IgA heavy chains had undergone somatic hypermutation, as had many of the IgM BCRs (Extended Data Fig. 2c). Samples were limitied, but overall the BCR repertoire observed at these late timepoints suggested that vaccination and/or parasite challenge did not promote generation or maintenance of a memory CSP-specific B cell population with significantly restricted HC usage.

### Generation and specificity of monoclonal antibodies

To address BCR specificity at a single-cell level, 24 (8 from each subject) of the 164 paired heavy and light chain sequences identified from CSP-specific B cells were selected for detailed analysis. We selected BCRs from the day of the second challenge (Timepoint C) and a later timepoint, 3 months later (Timepoint F), as these each represented memory timepoints. Again, we selected BCRs comprising a diverse array of phenotypes denoted by differential expression of cell surface proteins, HC and LC gene usage and mutation frequencies. In addition, we also expressed all IgA BCRs that were isolated (n=4). Further, to ensure that we were not excluding potential NANP-repeat-specific clones, we expressed all IgG BCRs expressing either VH3-33, VH3-30, or VH3- 23 from all timepoints (n=7.) In total, we expressed and screened 34 unique mAbs (Extended Data Table 3). All antibodies were again expressed with an IgG1 constant region regardless of their original isotype, and any IgM-derived antibody that didn’t bind as IgG1 was expressed as IgM.

This panel of recombinant mAbs was first screened for reactivity to full-length CSP by ELISA with 26 mAbs exhibiting CSP-specific binding (Fig. 2d, Extended Data Table 3). Next, we assessed IgG1 mAb reactivity to the NANP-repeat vs. C-term regions by peptide ELISA. Surprisingly, while a small proportion of mAbs recoginized the NANP-repeat region (4/17), the majority (13/17) were specific to the C-term region (Fig. 2d,e). C-terminal specificity was present in all subjects and included a range of VH genes within each subject: 3 for subject 71C, 5 for subject 71D, and 4 for subject 71E. In total, 10 different VH genes resulted in C-terminal specificity, including VH3-30 and VH3-23, previously associated with NANP-repeat-specificity (Extended Data Table 3). These findings were in direct contrast to our observations in different subjects from the same trial examined at an earlier timepoint (day 7 post dose 3) and who received a different vaccine regimen (3 standard doses, 1 month apart), (Fig 1), where mAbs were predominantly specific for NANP-repeat region. Together, these observations suggested a major shift in the circulating, CSP-specific B cell repertoire over time following repeat immunization and/or CHMI.

Based on these results, we further characterized 10 C-term-specific mAbs to begin to map C-term epitope binding activity more granularly. We performed a competitive ELISA using 2 previously identified C-term-specific antibodies (236 and 1512) known to bind to two distinct regions of the C-terminus of CSP^23^. All of the antibodies bound full-length CSP (Fig 3a). Four of the 10 antibodies blocked the binding of antibody 236 (Fig. 3b), suggesting they bind a similar region of the protein. Antibody 1512 was blocked strongly by 1 mAb and weakly by 2 more (Fig. 3c). Binding of 3 mAb was not impacted by either 236 or 1512, indicating that they likely targeted alternative C-term epitopes (Extended Data Table 4). These results suggested that a diverse population of B cells targeting at least 3 distinct epitopes within the CSP C-terminus comprise the memory B cell population at timepoints post-immunization and post-malarial challenge.

**Fig. 3:**
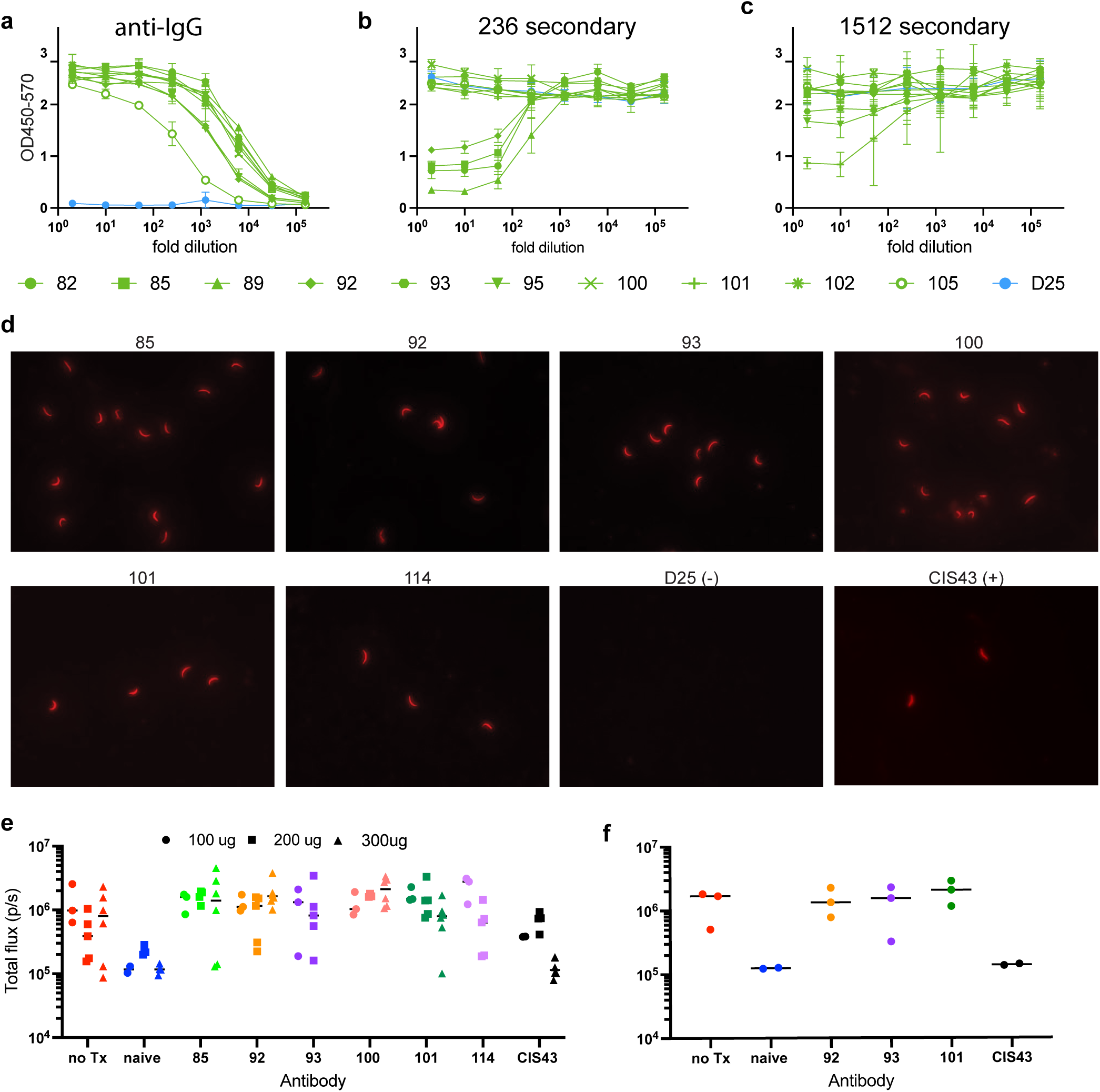
Epitope specificity, sporozoite binding and protective capacity of C-term-specific antibodies. a-c, Competitive ELISAs to determine epitope specificities of C-term mAbs All plates were coated with FL-CSP, incubated with C-term mAbs antibodies followed by HRP-conjugated **a,** anti-IgG, or two previously identified C-term-specific mAbs **b,** 236 or **c,** 1512 secondary. ELISA were conducted in triplicate and plots are representaive of 2 independent experiments. **d,** Binding of C-term-specific mAbs to sporozoites by immunofluorescence assay (IFA). Fluorescent microscopy images of C-term-specific mAbs bound to Pb-PfCSP-GFP-luc parasites are shown. RSV-specific antibody D25 was used as a negative control and CIS43 was used as a postive control. Data are representative of 5 independent experiments. **e,** In vivo protection from Pb-PfCSP-GFP-luc intravenous (i.v.) challenge 2 hours after administration of various doses of C-term specific antibodies. Parasite liver burden was analyzed by IVIS 42 hours post-infection. Known protective antibody CIS43 was used as a positive control. Data are representative of 3 independent experiments.**f,** In vivo protection from P.b-P.f.CSP-GFP-luc following pre-mixing antibodies with parasites prior to i.v. injection. Liver burden was determined as above. Murine CIS43 antibody was used as a positive control. This experiment was only conducted once.

### Functional characterization of C-term-specific antibodies

Given the long-term protection of these individuals and the profound shift of memory B cell specificity towards the CSP C-terminus region in all subjects tested at later timepoints, we next performed a detailed functional assessment of a panel of isolated C-term-specific mAbs. We selected 6 C-term-specific antibodies for detailed characterization based on varying HC usage and epitope specificity. We first measured binding affinities for full-length CSP and a C-terminus peptide (p16) by biolayer interferometry (BLI). Antibodies 236 and 1512 were included as positive controls for C-term binding, and NANP-repeat-binding antibody, CIS43, was included as a control for binding to FL-CSP. All 6 of the newly identified C-term mAbs bound both FL-CSP and p16 with relatively high affinities (1.3-68.2 nM) (Extended Data Table 4) comparable to all 3 control antibodies (FL-CSP: CIS43=3.3nM, 1512= 5.85nM, 236=17.9nM. p16: 1512=6.93nM, 236=0.97nM).

Next, we evaluated the capacity for these mAbs to bind CSP in the context of whole sporozoites. For this assay, we utilized sporozoites isolated from *Plasmodium berghei* previously engineered to express *P. falciparum* CSP in association with cis-linked GFP and luciferase (Pb-PfCSP-GFP- luc)^36^. In this system, expression of *P. falciparum*-derived CSP in the context of a rodent-specific parasite allows for *ex vivo* assessment of sporozoite binding as well as assessment of possible protection *in vivo* in an established mouse model of malarial infection. Whole sporozoites were isolated and fixed to slides and stained with individual C-term-specific antibodies followed by a fluorescently labeled anti-IgG antibody and imaged by microscopy. All 6 C-term mAbs exhibited sporozoite labeling, mirroring a NANP-repeat-binding positive control mAb (CIS43), while no labeling was observed for a RSV-specific negative control antibody (Fig. 3d).

To test the protective capacity of these mAbs in this mouse model, we employed passive monoclonal antibody transfer followed by IV infection using Pb-PfCSP-GFP-luc sporozoites. C57BL/6 mice were first treated with 100 μg antibody, a dose commonly used to assess protection in this model system,^37^ and infected 2 hours later with Pb-PfCSP-GFP-luc parasites. Forty-two hours later, parasite burden in the liver was assessed by quantifying luciferase activity by IVIS. Mice treated with each of the six mAbs showed no reduction in liver parasite burden compared to the non-treated, infected controls (Fig. 3e). Increasing mAb doses to 200 or 300 μg also failed to protect from infection as parasite levels in the liver were similar to the untreated group. In contrast, treatment with positive control antibody (CIS43) greatly decreased liver burden, especially at the 300μg dose. As these antibodies failed to protect even at high doses, we also tested protection in the setting of pre-mixing of mAb with sporozoites prior to injection into mice. This approach ensures that the lack of protection did not reflect a deficit in localization or mAb half-life relative to the control. The C-term mAbs also failed in this less stringent context (Fig. 3f). Collectively, these findings demonstrate that these 6 C-term-specific mAbs derived from CSP-specific B cells in vaccinated individuals do not protect in this murine model of infection. Future experiments are required to determine whether alternative mAbs, delivery methods and/or infection routes may result inprotection.

### Assessment of CSP-specific B cell responses over time in individual vaccinated subjects

Our data suggested the presence of a predominantly class-switched NANP-repeat specific memory B cell population at an acute timepoint shortly post-vaccination, but a more robust C-term-specific memory B population at late timepoints post-vaccination. Yet these studies were performed in different individuals, who had undergone different vaccination regimens, with relatively few cells analyzed. We therefore next wanted to determine whether these findings held true in the same individuals across the various timepoints. Because we had identified key CSP specificities to monitor and to remove any potential bias introduced in the selection of BCRs to be cloned and tested for specificity, we elected to employ a dual-tetramer approach to identify full-length CSP vs. C-term-specific B cells for single-cell analysis. We reasoned that any BCR that bound the full-length tetramer but did not bind the C-term tetramer was NANP-repeat-specific, as these are the only 2 regions included in RTS,S. To confirm this prediction and approach, we again sorted cells that bound full-length CSP and then sequenced and expressed 10 BCRs, five from cells that only bound the full-length tetramer and five from cells that also bound the C-term tetramer. These BCRs were selected to include a variety of VH and VL chain genes, including several VH3-33 clones (which is usally associated with repeat specificity) but which exhibited both binding to FL-CSP only and to the C-terminus tetramer (Extended Data Table 5). We then tested their specificity by ELISA. Nine of the 10 antibodies bound FL-CSP and of these, 4/4 that bound C-term tetramer only bound C-term peptide by ELISA and 5/5 that were C-term tetramer negative bound NANP-repeat peptide (Extended Data Table 5), validating this approach for subsequent analyses.

Using this dual-tetramer approach, CSP-specific B cells were isolated from samples taken at day 7 after the third vaccination (timepoint B*^7^) (Fig. 4a) for subjects 71C, 71D, and 71E (previously examined at memory timepoints on or after the second CHMI; Fig. 2 timepoints C-F). The majority of CSP-specific cells from all 3 subjects bound the full tetramer but exhibited limited C-term reactivity (23-26%), implying that the majority (73-76%) were NANP-repeat-specific as suggested by our prior data (Fig. 4b). When the phenotype of the cells was analyzed in more detail, both repeat- and C-term-specific plasmablasts were detected, comprising roughly 20% and 50% of the populations, respectively. Furthermore, of the non-plasmablasts, ∼1/2 of the NANP-repeat-specific B cells were CD21+CD27- naïve while ∼1/2 were memory cells (Fig. 4c), and of those memory cells, ∼ 1/2 were class-switched to IgG (Fig. 4d). Of the relatively smaller pool of non-plasmablast C-term-specific B cells, greater than 80% expressed memory markers (Fig. 4e), with nearly all of these expressing IgG (Fig. 4f). These data support our previous observation (Fig. 1) of a larger proportion of repeat-specific memory B cells relative to C-term at this acute timepoint post-third vaccination. Additionally, although there there were relatively fewer C-term-specific cells present, they appear to be phenotypically distinct from the repeat-specific cells, hinting at an altered memory cell differentiation program of the two epitope specific populations. Together, these findings support the conclusion that the skewing towards C-term specificity observed at memory timepoints reflects changes based on the time post-vaccination and/or CHMI and not differences unique to the subjects evaluated at each timepoint or the cells selected for B cell cloning.

**Fig. 4:**
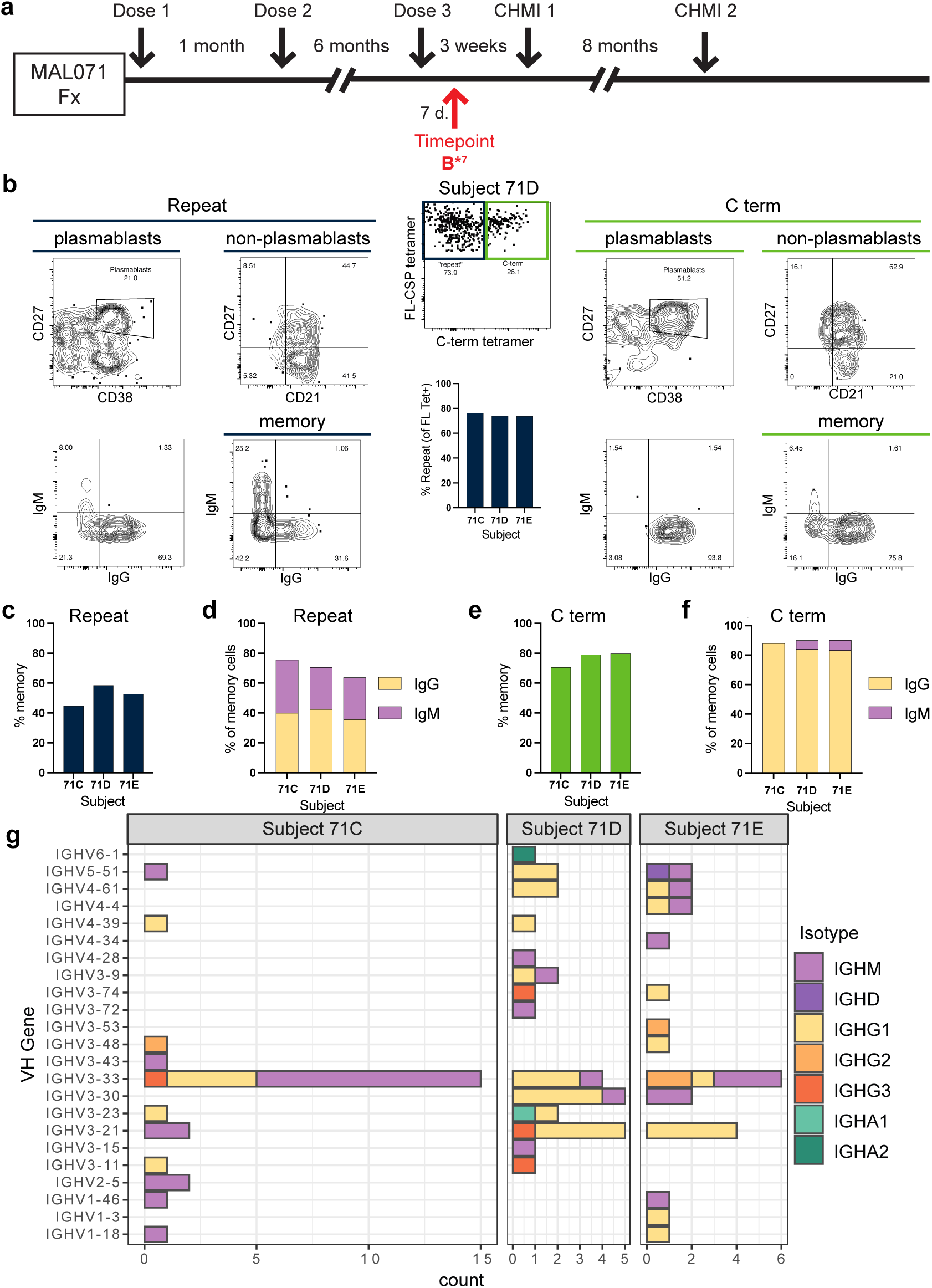
Repeat-specific B cells are more prevalent immediately following vaccination. **a**, Timeline of vaccination/challenge regime for MAL-071 Fx trial arm and PBMC sampling. PBMC samples analyzed seven days post-third dose (Timepoint B*7). **b,** Example flow plots of CSP-specific cells showing binding to a C-term specific tetramer. Repeat and C-term specific cells were further analyzed for suface expression of memory markers (CD27,CD21) and antibody isoypes. All gates were set on non-antigen specific popuations. Bar graph shows percentage of CSP tetramer+ cells that were repeat-specific for all 3 subjects. **c,** Percentage of repeat specific cells that exhibited memory phenotypes. These numbers include cononical memory cells (CD27^+^CD21^+^), activated memory (CD27^+^CD21^-^) and atypical memory (CD27^-^CD21^-^). **d,** Percentage of repeat specific memory cells expressing IgG or IgM. **e,** Percentage of C-term, specific cells that exhibited memory phenotypes. These include all memory phenotypes as described in **b**. **f,** Percentage of C-term specific memory cells expressing IgG or IgM. **g,** BCR heavy chain gene usage of single cell sorted CSP-specific B cells. The isotype indicated was determined by sequencing.

To evaluate the BCR gene usage and isotype of B cells with C-term vs. NANP-repeat-specificities at this early timepoint post-vaccination, a total of 252 CSP-specific cells were sorted. We obtained 92 pairs of sequences from the 3 subjects and examined V gene usage for both heavy and light chains (Fig. 4g). While broad VH usage was observed, VH3-33, VH3-30 and VH3-21 were predominant in all three subjects, with VH3-33 overrepresented in subject 71C, and most expressing the IgM isotype. Of note, at timepoint B*^7^, IgG cells expressing the NANP-repeat-associated heavy chains were present, but these were mostly lost by the memory timepoints (timpoints C-F) in these same individuals (Fig. 2c). Light chain usage was even more diverse, covering many different V genes and families. Interestingly, while the V gene usage was often similar between subjects, the pairing of light and heavy chains was diverse (Extended Data Fig. 3a).

The sequences of V genes were also examined for mutations from germline sequences (Extended Data Fig. 3b). All subjects had some mutations in their repertoires, although subject 71C did have a larger proportion of sequences without any mutations compared to the other subjects, and these were mostly mu isotype. The amount of mutations correlated with heavy chain isotype, with gamma and alpha having more mutations than mu for all subjects. Collectively, both the flow cytometric and sequencing data reveal the presence of repeat-specfic, class-switched memory B cells 7 days post-3^rd^ vaccination in the same individuals where these cells were largely absent at later times post-immunization (Fig. 2).

### Assessment of the MBC response in a separate vaccine trial

The proportional decline in NANP-repeat-specific memory B cells at late timepoints in a limited number of vaccinated subjects led us to extend our analysis to a larger number of individuals in an independent clinical trial, MAL092. This new study included both protected and non-protected individuals providing a possibility of identifying B cell phenotypes predictive of protection. Using our dual-tetramer approach, we examined PBMCs from 5 protected and 5 non-protected individuals at 3 memory timepoints: 1) 30 days following the third vaccination (Timepoint B*^30^); 2) the day of CHMI challenge (90 days post-last vaccination; Timepoint C*); and 3) 90 days post-challenge (180 days post-last vaccination; Timepoint F*) (Fig. 5a).

**Fig. 5:**
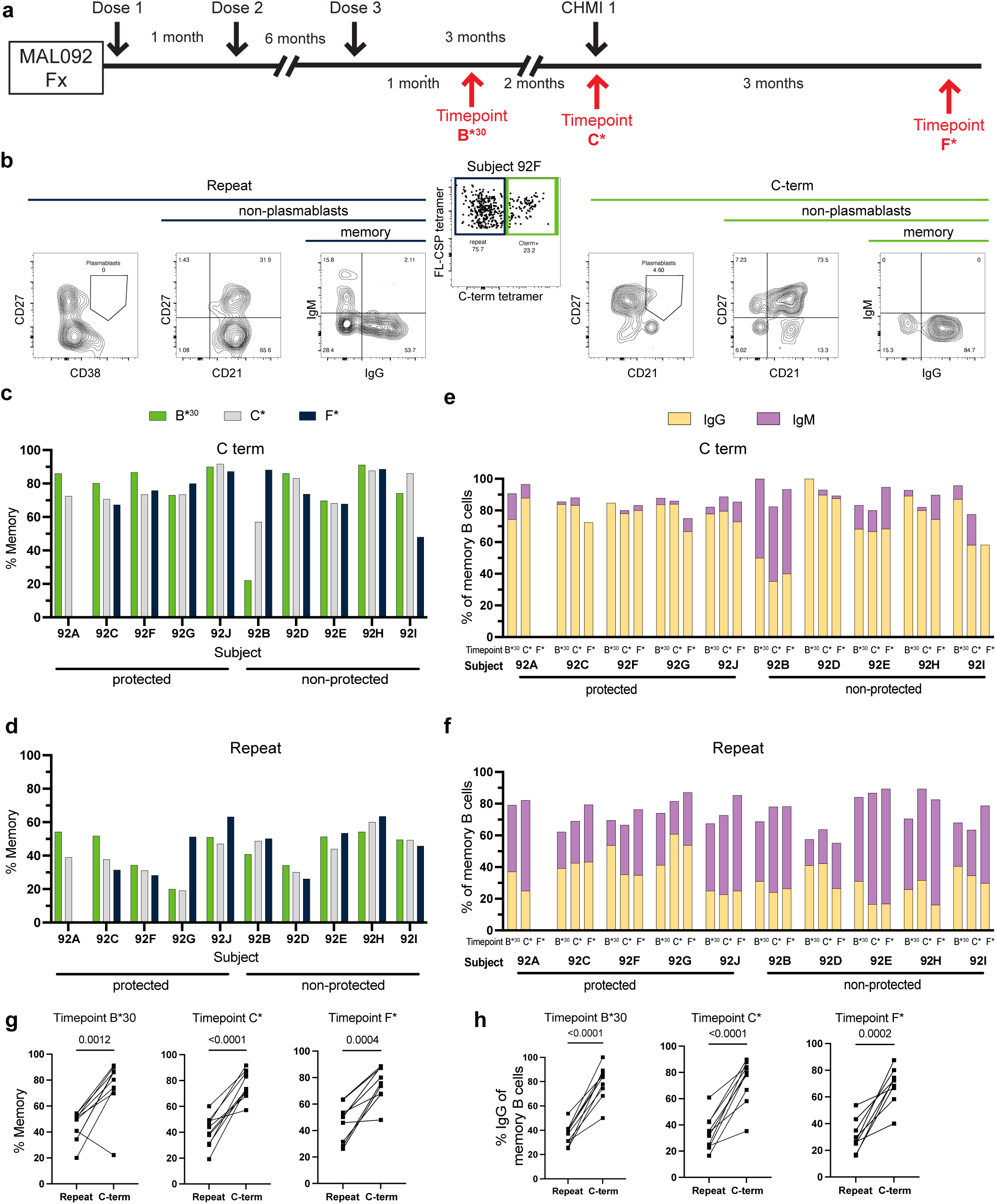
In both protected and unprotected individuals,the repeat-specific response is dominated by naive cells while the C-term specific response is dominated by IgG+ classical memory cells. **a**, Timeline of vaccination/challenge regime for MAL-092 Fx trial arm and PBMC sampling. PBMC samples were analyzed one month post dose 3 (Timepoint B*30), the day of second CHMI (Timepoint C*) and approximately three months post challenge (Timepoint F*). **b,** Example flow cytometry analysis of CSP-specific B cells from subject 079 at timepoint B*30. “Memory” includes all non-naive (CD21^+^CD27^-^) cells. Fraction of **c,** C-term or **d,** repeat region specific B cells expressing markers of memory cells. Fraction of **e,** C-term or **f,** repeat specific memory B cells expressing IgM or IgG. **g,** Comparison of the freqency of repeat- or C-term-specific cells expressing markers of memory in each study subject. **h,** Comparioson of the frequency of repeat- or C-term specific memory cells expressing IgG. Statistics were determined by 2-tailed Wilcoxon signed-rank tests.

Across these memory timepoints, C-term-tetramer binding cells only comprised 3-30% of circulating B cells (Fig. 5b, Extended Data Fig. 4a). While this overall proportion of C-term specific cells was lower than in the MAL071 subjects evaluated, the vast majority of the C-term-specific cells exhibited markers of memory (Fig. 5 b,c). In contrast, a much smaller proportion of the NANP-repeat-specific were memory, (CD27^+^CD21^+^, CD27+CD21^-^ or CD27^-^CD21^-^) with the majority being naïve (CD27^-^CD21^+^) (Fig. 5b,d). This skewing was further enhanced when looking at BCR isotype, with nearly all the C-term-specific memory cells being class-switched to IgG (Fig. 5e), while on average fewer than 50% of the NANP-repeat-specific memory cells expressed class-switched BCRs (Fig. 5f). When the relative frequencies (across all subjects at each timepoint) were compared between NANP-repeat- vs. C-term-specific B cells, C-term-specific cells were significanlty enriched for both memory phenotype (Fig. 5g) and IgG expression within memory cells (Fig. 5h). Thus, although the majority of the cells present in these individuals at these memory timepoints are repeat-specific, relatively few were class-swiched memory cells. Collectively, these results align with our previous findings that the proportion of C-term-specific MBCs increase with time since the 3^rd^ vaccination, while repeat-specific MBCs decline. Comparing B cell features in individuals over time, no major differences in B cell phenotypes (Fig. 5b-f) or specificity (Extended Data Fig. 4a) were observed. This suggests that the memory response was established within ∼30 days post-vaccination and minimally altered by parasite challenge. We also did not observe any obvious differences between protected and non-protected individuals.

We next selected 2 subjects from each of the protected (subject 92C, 92F) and non-protected groups (subjects 92D,92E) for BCR sequencing at the pre-CHMI memory timepoint (Timepoint C*) (Fig. 5a). Both individuals displayed a wide range of VH and VL gene usage. VH3-33, the only over-represented HC, was overwhelmingly derived from NANP-repeat-specific B cells (Fig. 6a and Extended Data Fig. 4b). Sequencing confirmed that the majority of the C-term-specific cells were class-switched while the NANP-repeat-specific cells were not, and nearly all of the class-switched NANP-repeat-specific cells utilized the VH3-33 heavy chain (Fig. 6b and Extended Data Fig. 4c). We also evaluated the VH mutation frequency, demonstrating that the majority of C-term-specific receptors had undergone somatic hypermutation, and the few lacking mutations were IgM derived. In contrast, the proportion of NANP-repeat-specific BCRs with mutations was much lower, with only the few class-switched cells and a handful of IgM cells having been mutated (Fig. 6c). This may reflect the fact that many repeat-specific heavy chains likely require less mutation to become high affinity due to the higher baseline affinity of the germline sequene for the repeat region. Overall, these results align with what we observed in the MAL071 trial subjects, supporting the conclusion that RTS,S/AS01 vaccination results in a limited NANP-repeat-specific memory response and promotes the generation of a MBC pool that is increasingly comprised of class-switched C-term reactive cells.

**Fig. 6:**
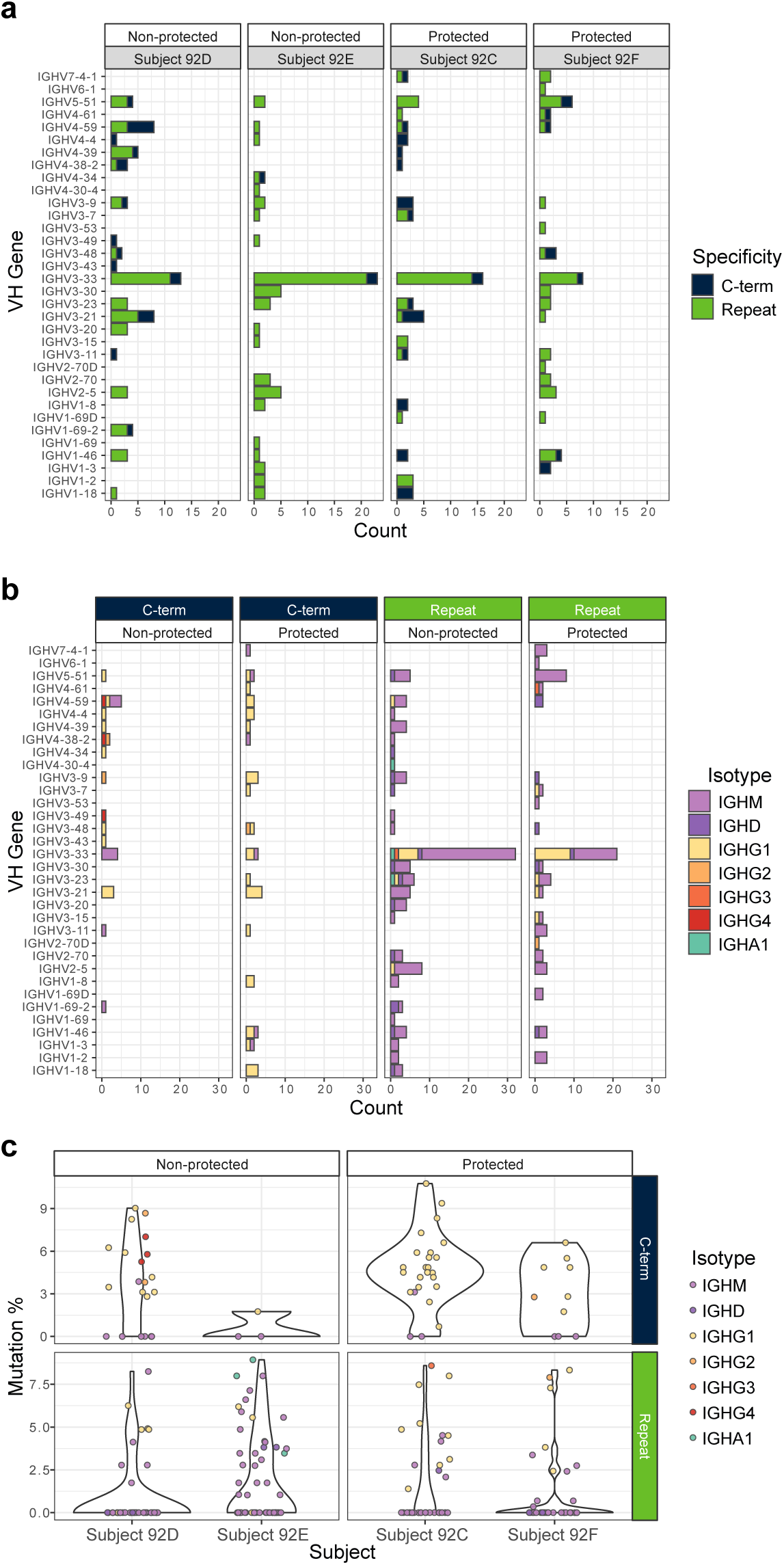
Heavy chain gene usage and somatic hypermutation rate of C-term vs Repeat BCRs. **a**, BCR heavy chain gene usage of single CSP-specific B cells sorted from 2 protected and 2 non-protected individuals. The specificity indicated was determined by tetramer binding to the cells during sorting. **b,** Heavy chain gene usage broken down by isotype BCR isotype. Data from the 2 protected and 2 non-protected subjects has been combined. **c,** Somatic hypermutation rates of sequenced BCRs.

## Discussion

RTS,S has proven to be protective in numerous clinical trials. However, protection mediated by the vaccine appears to be short lived in most individuals^5,32^. While extensive efforts have be made to determine which aspects of the immune response mediate this protection^17,32,38–41^, precise mechanisms of protection have not been identified. One of the strongest associations with protection identified to date is the production of antibodies against CSP, and in particular the repeat region of this protein^9^. Additional support for this concept has been derived from the development of mAbs that bind junctional and/or repeat epitopes and exhibit robust protective capacity in both mice and humans^2,4,28,42–44^. While we did not evaluate junctional-specific responses, our results demonstrate that class-switched NANP-repeat-specific B cells dominate the response early after immunization (Figs. 1 and 4) and contain a significant proportion of memory B cells (Fig. 1). Strikingly, however, while NANP-repeat-specific B cells remain detectable, and were predominant in subjects studied from the MAL092 cohort, memory B cells with these specificities decline in the blood (Fig. 3), and were reduced in frequency by 30 days post-immunization (Fig. 5). These results were even more profound when looking at IgG class-switched cells. These findings correlate with studies that show a relatively rapid decrease in anti-NANP-repeat-specific antibody titers post-vaccination^5–7^. Given that both memory cells and plasma cells can be derived through similar germinal center-dependent processes, these observations suggest that the vaccine may lead to a limited pool of long-lived plasma cells (PC), which is supported by diminishing antibody titers in prior studies, and that boosting of the NANP-repeat-specific antibody response following re-immunization may be limited by the diminished memory cell pool capable of responding to the boost^9,11,24,45^. Taken together, our results in association with previous reported findings suggest that vaccine-generated, NANP-repeat-specific B cells primarily generate short-lived antibody secreting plasmablasts and exhibit limited capacity to generate long-lived MBCs that are able to respond to booster doses or infection to increase protective antibody levels. More extensive studies examining B cell phenotype and specificity thoughout the course of immunization and challenge are needed to confirm these hypotheses.

It has been reported that suppressive antibody feedback may be responsible for altering CSP-specific responses following repeated vaccination^46^. In these studies, it was demonstrated that increased levels of NANP-repeat-specific antibodies dampen subsequent NANP-repeat-specific B cell responses while driving expansion of the C-term-specific cells. These results generally align with ours and may be one mechanism driving the shift in specificity away from the NANP-repeat region and towards C-term at the memory timepoints. However, since we did not examine the response any earlier than post-third vaccination, combined with the vastly different vaccine and and vaccination schedules used in these studies, direct comparison with our results is challenging. It has also been suggested that delaying the final booster may overcome some of the effects of suppressive antibody feedback and increase the ability to boost CSP-based vaccines^46^. However we observed limited numbers of class-switched, somatically mutated NANP-repeat-specific memory cells 30 days after the delayed fractional dose (Figs. 5 and 6), suggesting that memory B cells with NANP-repeat specificity either did not become reactivated and expanded and/or did not undergo a germinal center response.

While few NANP-repeat-specific IgG-expressing MBCs were detected by day 30 post-third immunization in our studies, B cells specific to the C-term of CSP were more readily abundant and included class-switched, affinity-matured MBCs. These C-term-specific MBCs persisted at all subsequent timepoints examined (up to approximately 1 year following third vaccine dose) (Figs. 2, 5, 6). The role of C-term antibodies in RTS,S-mediated protection remains unclear^15,40,41,47^. C- term antibody avidity^15^ and Fc region mediated effector fucntions^40,41^ have been shown to correlate with protection. However, only one C-term-specific mAb has shown some activity in mouse models, at levels of protection much below that of repeat/junctional region targeting antibodies^23^. Most C-term antibodies show no activity in these models^21,22^. We generated a panel of C-term-specific antibodies from B cells isolated at memory timepoints including mAbs that bound CSP with high affinity (Fig. 2, Extended Data Table 4). However, despite assessment of a diverse pool of C-term-specific antibodies isolated from germinal center-derived B cells, we failed to identify protective activity in an intravenous murine challenge model. The lack of protection conferred by C term mAbs in this study (Fig. 2, Extended Data Table 4) suggests that the long term protection observed in the individuals examined was not due to the C-term dominant B cell populations they possessed at the memory timepoint, but perhaps from either elicitation of repeat-specific antibodies maintained at sufficient levels for protection or a non-humoral immune mechanism, such as T cells in the liver. However, it remains possible that C-term antibodies might mediate protection in alternative challenge models; and further study will be needed to confirm the lack of protective capacity.

While we observed a predominance of MBCs in the C-term specific pool at all memory timepoints in all subjects across both vaccine cohorts, NANP-repeat-specific IgG-expressing MBCs were present, albeit at a lower frequency than at earlier timepoints. The reason for the diminished ability of NANP-repeat-specific cells to persist long-term after vaccination was not addressed in this study, but is addressed in an accompanying manuscript (McDougal et al.). In accordance with data from our concurrent study, we hypothesize that loss of protective NANP-repeat-specific memory may be due to the increased avidity associated with the repeat epitope. Highly repetitive antigens such as proteins with repeated motifs and polysaccharides have been shown to preferentially drive the formation of short-lived plasmablasts^18,48–55^. These responses are often T-independent, but it has been shown that the repeat-specific CSP B cell repsonse does require T cell help^56^, but the high avidity of the repeat-antibody interaction may still skew B cells to a short- lived plamablast fate. We hypothesize that this skewing to short-lived ASCs production also occurs following RTS,S immunization, with the NANP-repeat-specific B cells differentiating into antibody-secreting plasmablasts capable of mediating protection in most individuals until antibody titers drop below protective levels. In contrast, our combined findings suggest that the non-repetitive C-term epitopes within the vaccine are able to generate long-lived memory B cells; however, these specificities appear not to be protective. Additionally, the clinical trials used for our current study were conducted in malaria-naïve adults, which is not target population of this vaccine. McDougal et al offers insights into differences in repeat and C-term specific B cell responses to vaccination in both adults and children in malaria endemic settings.

Current vaccine strategies aimed at improving upon the success of RTS,S by altering adjuvants, vaccine schedules or copy number of antigen^57^ may suffer similar consequences by eliciting responses analogous to that observed in our study as they share the same core antigen. Inclusion of a high copy number of repeat-containing antigen is predicted to preferentially stimulate B cells specific to this region to a short-lived, terminally differentiated, plasmablast response. While these offer short-term protection, alternative approaches that include a lower total copy number of NANP-repeats, less repetitive regions of CSP (such as the junctional region) and/or antigens other than CSP might yield better long-term protection by inducing long-lived plasma cells and MBC populations required to maintain protective antibody titers.

## Materials and Methods

### PBMC samples

Frozen human PBMC from the clinical trials NCT01857869 (MAL071) and NCT03162614 (MAL092) were provided by PATH and GSK (study sponsor). Samples were thawed in PBS contain 10% FBS then washed and resuspended in complete media.

### Tetramer generation

Recombinant biotinylated full-length CSP and C-terminus peptide (p16) were provided by PATH. The proteins were tetramerized with streptavidin-PE or streptavidin-APC (Agilent), concentrated to 1 µM and stored in 50% glycerol at -20°C as previously described^26^. Decoy reagent was generated as previously described^58^.

### Tetramer staining and sorting

PBMC were stained with decoy reagent followed by CSP tetramers prior to incubation with anti-PE magnetic beads and column enrichment as previously described^26^. Enriched cells were stained with surface antibodies for 20 minutes on ice. Cells were then index sorted on a FACSAriaII (BD) and single cells were collected in 96 well PCR plates containing SMART-Seq v4 capture buffer (Takara Bio).

### ELISA

2 µg/ml of recombinant CSP (rCSP), NANP-repeat or C-term peptide in PBS was added to high binding 96 well plates (Corning) and incubated overnight at 4°C. Protein coated plates were washed with PBS-T (0.05% Tween-20 in PBS) and blocked with PBS-T containing 3% milk for 1 hour at room temperature (RT). Purified mAbs or culture supernatants were serially diluted in PBS-T containing 1% milk, added to the plates in triplicate and incubated for 2 hours at RT. After washing, HRP conjugated anti-human IgG detection antibody (Jackson ImmunoResearch) was added at 1:3000 in PBS-T containing 1% milk and incubated for 1 hour at RT. Detection was done using 1X TMB Substrate Solution (Invitrogen), quenched with 1M HCl and optical density (450nm- 570nm) was measured on a spectrophotometer.

### BCR sequencing

MAL071 SD Timepoint A samples were sequenced following protocols previously described in detail^59^. All other samples used SMART-Seq v4 at half reaction volumes to generate cDNA from singly sorted B cells. For MAL-071 Fx Timepoints C-F, a Sanger sequencing method was used as previously described in detail^58,60,61^. For all other samples, an Illumina sequencing method was used (manuscript in preparation as Thouvenel et al). Briefly, a multiplex PCR was performed on cDNA from each B cell using pooled constant region primers for IgM, IgG, IgA, IgD, IgK, and IgL constant regions and well-specific barcoded forward primers for the universal template switch region. Half plates were then pooled and library prepped using enzymatic fragmentation and ligation with an anchored multiplex PCR approach^62^ to retain well-specific barcodes. Indexed library preps were run on an Illumina Miseq and reassembled using a custom script and 10X Genomics Cell Ranger VDJ 6.1.2. Additional VDJ annotations and mutation counts were performed with IMGT/HighV-Quest^63^ (GENE-DB Version 3.1.41)

### BCR cloning

Primers corresponding to the V and J sequences were used with first round PCR products to facilitate cloning. Each light chain was cloned into vectors with their respective constant regions for IgK, and IgL. Heavy chains were cloned into IgG1 vectors and a portion of IgM derived sequences were also cloned into IgM expression vectors. For MAL-071 SD Timepoint A, a restriction enzyme cloning method was used as described in detail previously^64^. For all other timepoints an in-fusion cloning kit (Takara Bio) was used. All plasmids were confirmed to match their original cDNA sequence by Sanger sequencing.

### Competitive ELISA

Plates were coated with full-length rCSP as described above. C-term-specific antibodies generated in these studies were diluted 2 fold followed by 7 5-fold serial dilutions. Antibodies were added to washed and blocked plates and incubated for 2 hours at room temp. Reference C-term-specific antibodies 236 and 1512 (provided by PATH) were conjugated to HRP using Lightning-Link HRP Conjugation Kit (Abcam) according to the manufacturer’s instructions. Plates were washed and 50ul of 10ug/ml 236 or 1512 were add and incubated for 1 hour at RT. Plates were then washed and developed as described above.

### Sporozoite-binding immunofluorescence assay (IFA)

Sporozoites (SPZ) were fixed in 500 ul 4% paraformaldehyde for 15-60 minutes at room temperature, spun at 8000xg for 2 minutes and most of the fixative was removed. SPZ were resuspended in approximately 60ul PBS and 5-10ul was spotted on to PTFE 12 well slides (Electron Microscopy Sciences) to achieve a few thousand SPZ per well. Slides were air dried in a fume hood until all liquid evaporated. SPZ were permeabilized in 20 ul 0.5% Triton X-100 for 30 minutes at RT in a humidity chamber, washed once with PBS and blocked with 3% BSA in PBS for 30 minutes at 37 C or 1 hour at RT. Following 3 PBS washes, C-term antibodies were added at a 1:100 dilution and incubated at 37 C for 1 hour or 4 C overnight. Wells were washed 3X in PBS and anti-GFP-FITC , anti-human IgG-Dylight594 and DAPI (Invitrogen) were added at 1:500 and incubated at 37C for 30 minutes. After 3 PBS washed, antifade reagent ProLong Gold (Invitrogen) was added and the slides were cover slipped before imaging.

### *In vivo* protection

Male C57BL/6 mice (The Jackson Laboratories) were given 100-300 µg anti-C-term or positive control CIS43 antibodies intravenously (IV) 2 hours prior to infection. 2000 freshly harvested *P. berghei* expressing *P. falciparum* CSP, GFP and luciferase (Pb-PfCSP-GFP-luc)^36^ were injected IV. For antibody pre-treatment, antibodies and parasites were mixed and incubated on ice for 15 minutes prior to IV injection. 42 hours post infection, mice were treated with luciferin and after 5 minutes, liver burden was measured by IVIS Spectrum (Revvity). All mouse experiments were ethically performed in accordance with the University of Washington Institutional Care and Use Committee guidelines.

### Affinity measurement by biolayer interferometry (BLI)

BLI assays for affinity were performed at ambient temperature with shaking set at 1,000 rpm using an Octet Red 96 System (Pall FortéBio/Sartorius). Antibodies were diluted 5 µg/ml to in kinetics buffer (1× Hepes-EP+ [Pall FortéBio], 0.05% nonfat milk, and 0.02% sodium azide). Biosensors were hydrated in kinetics buffer for 10 min, and the diluted antibodies was immobilized onto Protein A biosensors (Greiner Bio-One) and then equilibrated in kinetics buffer for 60 s. Full-length CSP of C-term peptide was diluted to 500 nM in kinetics buffer and serially diluted threefold for a final concentration of 0.69 nM in a black 96-well microplate (Greiner Bio-One) at 200 μl per well. To measure association, loaded biosensors were dipped into the diluted protein and association and dissociation was performed for 60 s each. The data were baseline subtracted and plotted using Pall FortéBio/Sartorius analysis software (v12.0).

### Quantification and Statistical Analysis

Statistics are described in the figure legends and were determined using Prisim (Graphpad). All measurements within a group are from distinct samples, except for technical replicates used in ELISAs.Raw p values are reported. Raw sequencing data is available at XXXX (will added this once we have it deposited.)

## Acknowledgements

This study was funded by PATH’s Center for Vaccine Innovation and Access through grants from the Gates Foundation (OPP1108403 and OPP1191923/INV007217). We thank PATH’s Center for Vaccine Innovation and Access (CVIA) and GSK (Belgium) for their collaboration in this study and for providing PBMC samples. We thank the Walter Reed Army Institute of Research (WRAIR), MAL071 clinical trial PI Jason Regules, MAL092 clinical trial PI James Moon, GSK, and GSK staff (Johan Vekemans, Opokua Ofori-Anyinam) for conducting the clinical trials. We also thank the MAL071 and MAL092 vaccine trial participants and clinical staff. We thank Ulrike Wille-Reece for her role in conceptualizing and supporting this project and Cynthia Lee, Opokua Ofori-Anyinam, Katie Ewer, Ramiya Ravindranath and Bart Cuypers for their careful review of the manuscript.

---

**Extended Data Fig. 1:**
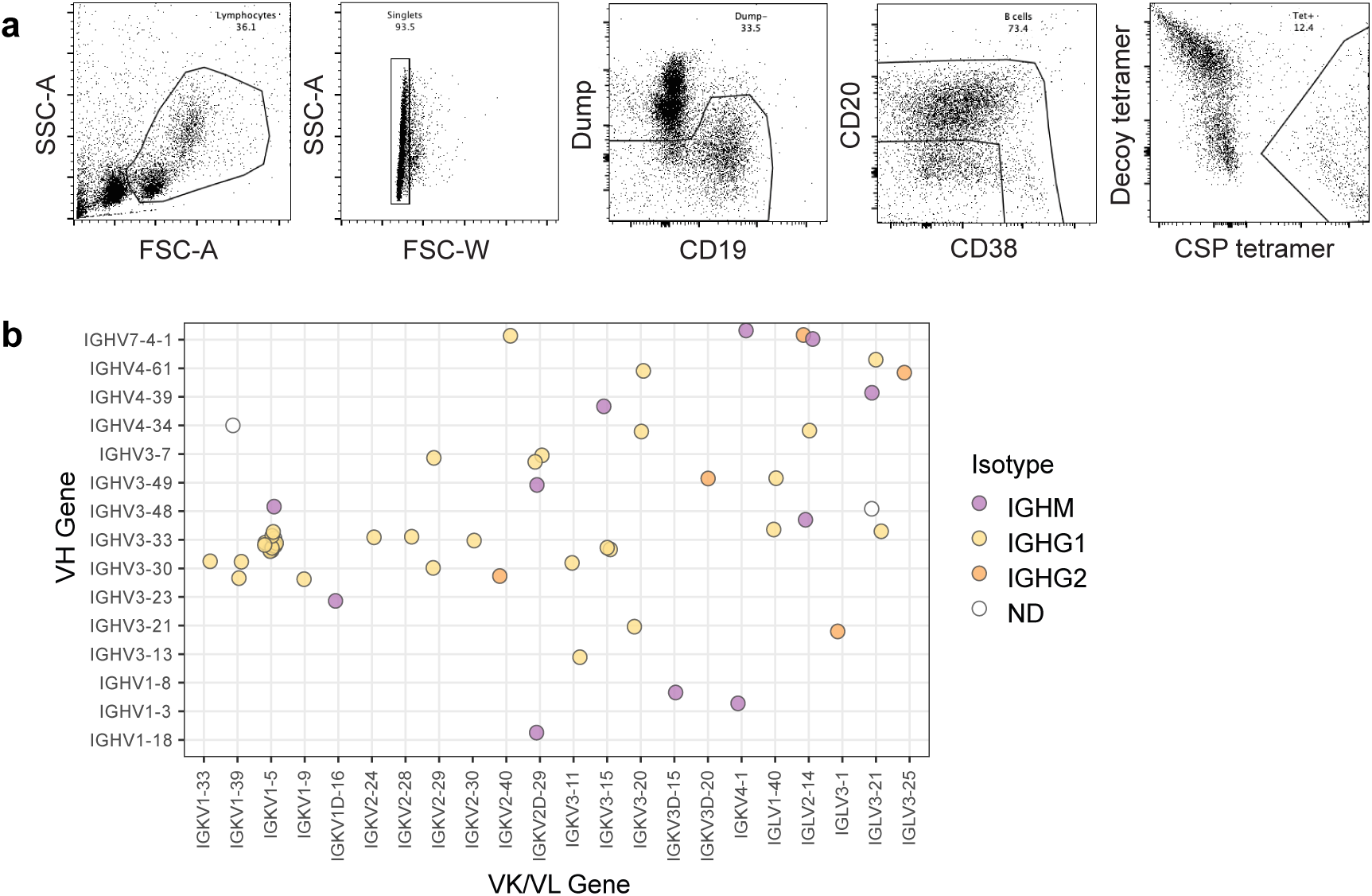
Sort gating strategy and additional BCR sequence analysis from MAL071-SD subjects 7 days post 3^rd^ vaccination. **a**, Example gating strategy used to sort single CSP-specific B cells from MAL071 trial subjects. **b,** Paired heavy and light chain gene usage for same cells shown in Fig. 1d. Data shown are a combination of data from subjects 71A and 71B.

**Extended Data Fig. 2:**
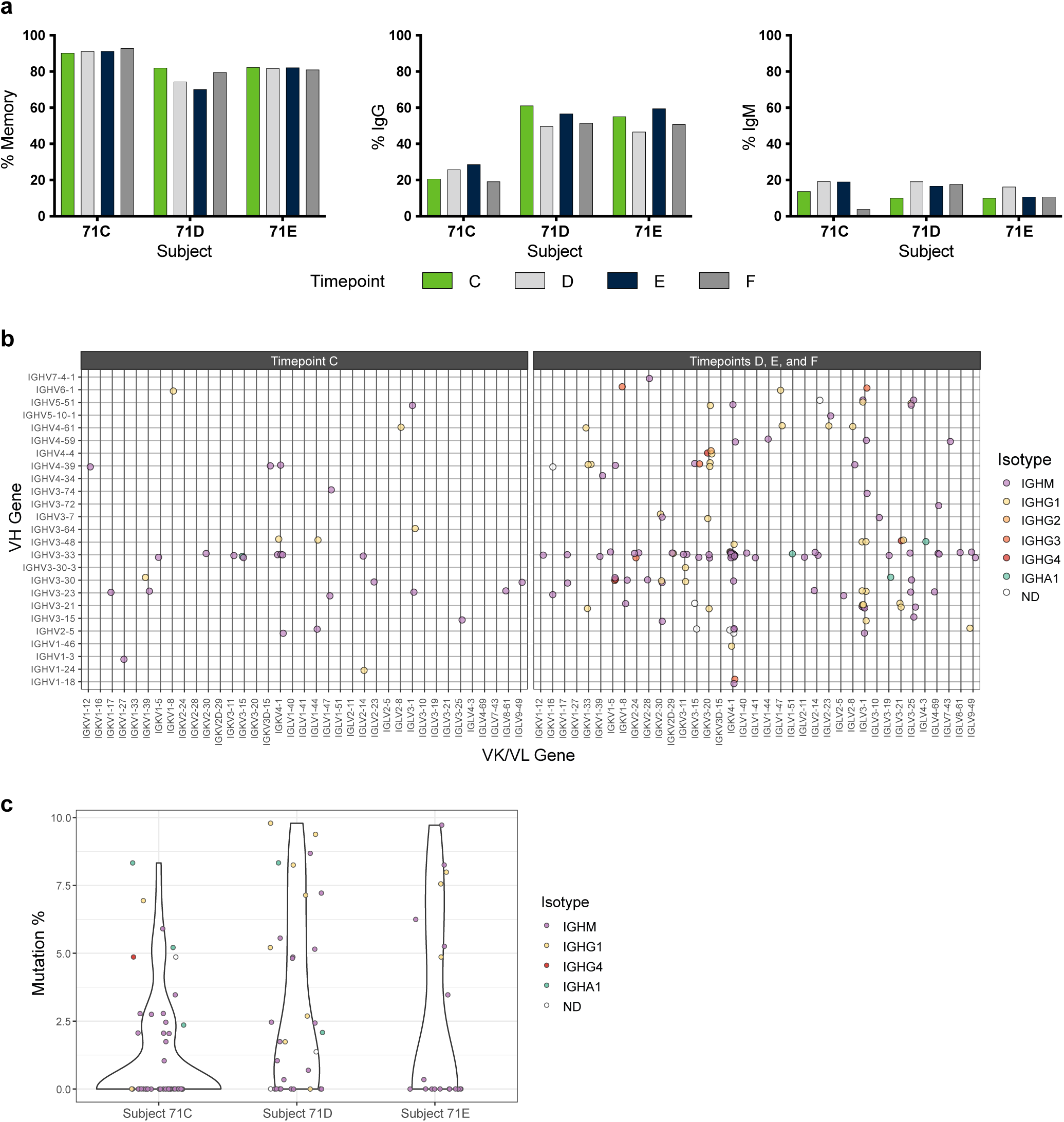
Additional flow phenotyping and BCR sequence analysis of MAL071-Fx subjects at day of second CHMI and beyond. **a**, Proportion of non-plasmablast cells exhibiting markers of memory, IgG and IgM. **b**, Paired heavy and light chain gene usage from all cells sorted at all timepoints. All subjects have been combined. **c,** Somatic hypermutation rates of sequenced BCRs.

**Extended Data Fig. 3:**
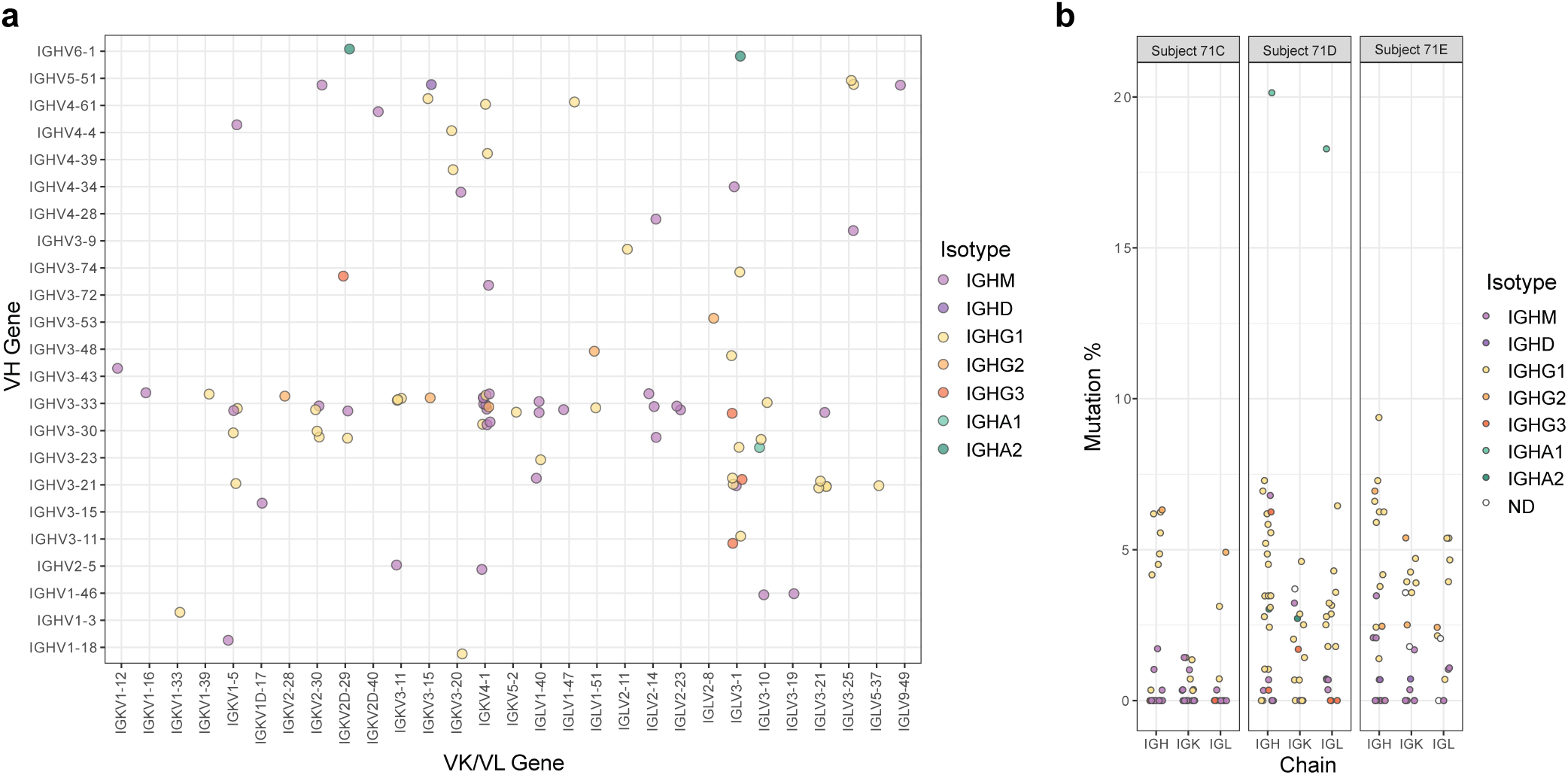
Additional BCR sequence analysis of MAL071-Fx subjects at day 7 post 3^rd^ vaccination. **a**, Paired heavy and light chain gene usage for cells shown in Fig. 4C. **b,** Somatic hypermutation rates by chain and isotype.

**Extended Data Fig.4:**
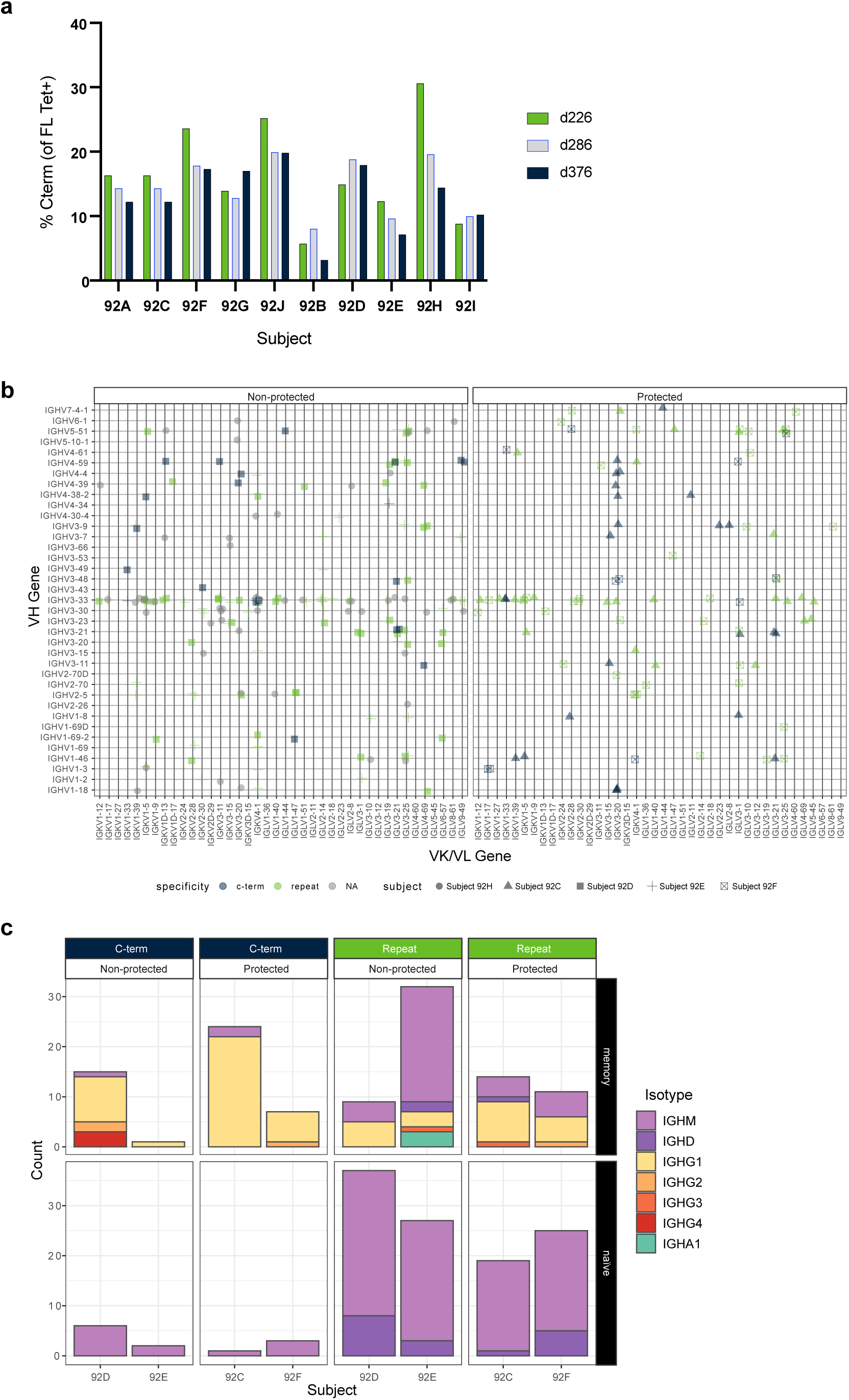
Additional flow phenotyping and BCR sequence analysis of MAL092-Fx subjects. **a**, Percentage of CSP tetramer cells that were C-term-specific. **b,** Paired heavy and light chain gene usage for the same cells shown in Fig. 6a. **c,** Total counts of cell expressing each BCR isotype for cells shown in Fig. 6b.

**Extended Data Table 1:**
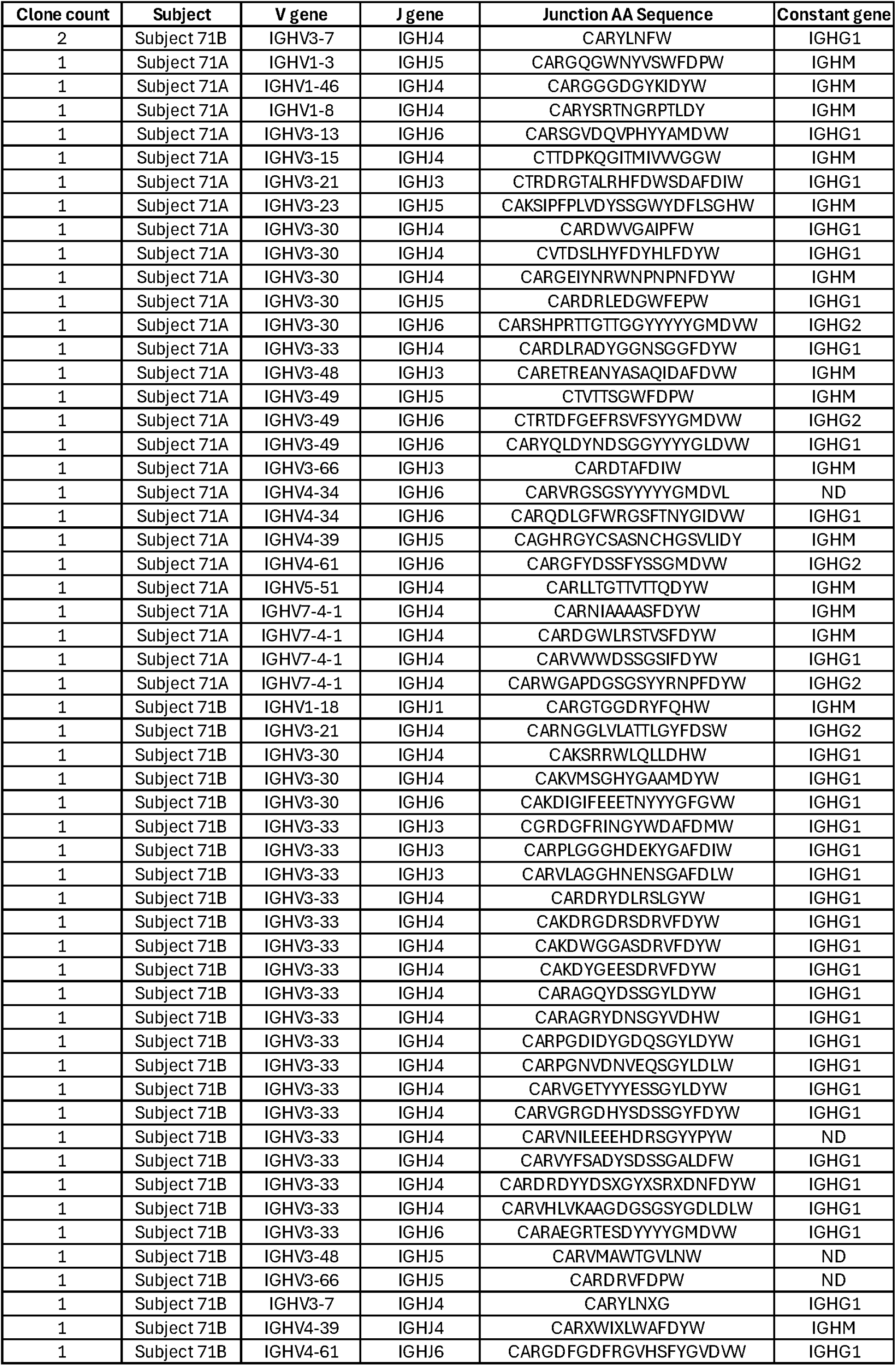
Clonality of sequenced BCR from Figure 1.

**Extended Data Table 2:** Clonality of sequenced BCR from Figure 2.

| clone count | Subject | V gene | J gene | Junction Sequence | Constant gene | Timepoint |
| --- | --- | --- | --- | --- | --- | --- |
| 2 | Subject 71D | IGHV3-23 | IGHJ4 | CANPWKDGYYGDSW | IGHM | C |
| 1 | Subject 71C | IGHV1-2 | IGHJ4 | CARDQGYDFWSGADYW | IGHM | E |
| 1 | Subject 71C | IGHV2-5 | IGHJ1 | CAHSSWGSYHHW | IGHM | C |
| 1 | Subject 71C | IGHV2-5 | IGHJ4 | CAHRHSAVTAAGTFDYW | IGHM | D |
| 1 | Subject 71C | IGHV2-5 | IGHJ4 | CAHSGSYYDFWSGSHPAFDYW | IGHM | C |
| 1 | Subject 71C | IGHV3-21 | IGHJ4 | CARESIAGVAKHFDYW | IGHM | F |
| 1 | Subject 71C | IGHV3-21 | IGHJ4 | CVIVILISTISYYMDYW | ND | F |
| 1 | Subject 71C | IGHV3-23 | IGHJ4 | CAKCLTGYYSIDYW | IGHM | D |
| 1 | Subject 71C | IGHV3-30 | IGHJ4 | CARDLWYRGDFDYW | IGHM | D |
| 1 | Subject 71C | IGHV3-30 | IGHJ4 | CAGGGYDYGDYVCLYW | IGHM | D |
| 1 | Subject 71C | IGHV3-33 | IGHJ1 | CARDLGSSGW | IGHM | D |
| 1 | Subject 71C | IGHV3-33 | IGHJ1 | CARDEGDNYFQHW | IGHM | D |
| 1 | Subject 71C | IGHV3-33 | IGHJ3 | CAATIPDAFDIW | IGHM | F |
| 1 | Subject 71C | IGHV3-33 | IGHJ3 | CAGGTGKGAFDIW | IGHM | D |
| 1 | Subject 71C | IGHV3-33 | IGHJ3 | CARDEDANGVYGW | IGHM | F |
| 1 | Subject 71C | IGHV3-33 | IGHJ3 | CAREGSIHAFDIW | IGHM | D |
| 1 | Subject 71C | IGHV3-33 | IGHJ3 | CALGYSSSLAFDIW | IGHM | D |
| 1 | Subject 71C | IGHV3-33 | IGHJ3 | CARPKVGEHAFDIW | IGHM | C |
| 1 | Subject 71C | IGHV3-33 | IGHJ3 | CASCLQELLHAFDIW | IGHM | F |
| 1 | Subject 71C | IGHV3-33 | IGHJ4 | CARDPIAAAPSW | IGHM | F |
| 1 | Subject 71C | IGHV3-33 | IGHJ4 | CARAAYSSSFDYW | IGHM | D |
| 1 | Subject 71C | IGHV3-33 | IGHJ4 | CARALIGGGITGYW | IGHM | D |
| 1 | Subject 71C | IGHV3-33 | IGHJ4 | CARDDSYSAFGGYW | IGHM | F |
| 1 | Subject 71C | IGHV3-33 | IGHJ4 | CARSGYSGSYFDYW | IGHM | C |
| 1 | Subject 71C | IGHV3-33 | IGHJ4 | CARDPFSGGDYVGYW | IGHM | F |
| 1 | Subject 71C | IGHV3-33 | IGHJ4 | CARDLGSVSGGSAEYW | IGHM | D |
| 1 | Subject 71C | IGHV3-33 | IGHJ4 | CARDLGWGSGWDLGYW | IGHM | C |
| 1 | Subject 71C | IGHV3-33 | IGHJ4 | CARESGYSSSRGFDYW | IGHM | D |
| 1 | Subject 71C | IGHV3-33 | IGHJ4 | CARVRDSSSSFLVDYW | IGHG1 | F |
| 1 | Subject 71C | IGHV3-33 | IGHJ4 | CARAGGRYSSGWCOLDYW | IGHM | F |
| 1 | Subject 71C | IGHV3-33 | IGHJ4 | CARPYGYDYGGYFDYW | IGHM | F |
| 1 | Subject 71C | IGHV3-33 | IGHJ4 | CARVGGYDFWWSGYFDYW | IGHM | D |
| 1 | Subject 71C | IGHV3-33 | IGHJ4 | CARDFGCFNGVCSNYLDYW | IGHM | E |
| 1 | Subject 71C | IGHV3-33 | IGHJ4 | CARDGSGSYGTSLEGGDYW | IGHM | F |
| 1 | Subject 71C | IGHV3-33 | IGHJ4 | CARDPGAGSSGPTPGGDYW | IGHM | F |
| 1 | Subject 71C | IGHV3-33 | IGHJ4 | CARAGDPRMYYYDSSSGSYGFDYW | IGHM | D |
| 1 | Subject 71C | IGHV3-33 | IGHJ5 | CARDLGSW | IGHM | D |
| 1 | Subject 71C | IGHV3-33 | IGHJ5 | CARHSSNYGFDNW | IGHM | D |
| 1 | Subject 71C | IGHV3-33 | IGHJ5 | CARDYSYQNWFDPW | IGHA1 | F |
| 1 | Subject 71C | IGHV3-33 | IGHJ6 | CAREGGAAAVGYMDW | IGHA1 | C |
| 1 | Subject 71C | IGHV3-33 | IGHJ6 | CARAGHDFWGLGMDW | IGHM | D |
| 1 | Subject 71C | IGHV3-33 | IGHJ6 | CARDGGVTADYYMYMDW | IGHM | E |
| 1 | Subject 71C | IGHV3-48 | IGHJ4 | CARAGVYSTSSGPTIPYW | IGHA1 | D |
| 1 | Subject 71C | IGHV3-48 | IGHJ4 | CARGWEDLPVAGFGPDPYFDSW | IGHG1 | C |
| 1 | Subject 71C | IGHV3-48 | IGHJ5 | CARLGIGFVWINNWFDPW | IGHG4 | F |
| 1 | Subject 71C | IGHV3-7 | IGHJ4 | CASLLLYDSSGYDWGYFDYW | IGHM | E |
| 1 | Subject 71C | IGHV3-72 | IGHJ3 | CARDGRGSSGTGDAFDIW | IGHM | D |
| 1 | Subject 71C | IGHV4-59 | IGHJ3 | CARSGGSSRDAFDIW | IGHM | F |
| 1 | Subject 71D | IGHV1-24 | IGHJ6 | XATNAMKGRYLDLFYPMDW | IGHG1 | C |
| 1 | Subject 71D | IGHV1-3 | IGHJ5 | CARLPFDPW | IGHM | C |
| 1 | Subject 71D | IGHV2-5 | IGHJ4 | CAHRSPCSRGSCSFEYR | ND | F |
| 1 | Subject 71D | IGHV3-15 | IGHJ4 | CTTGINW | IGHM | E |
| 1 | Subject 71D | IGHV3-15 | IGHJ4 | CTTVGAGTFDYW | IGHM | D |
| 1 | Subject 71D | IGHV3-21 | IGHJ4 | CARERRVAAPGYIDIW | IGHG1 | F |
| 1 | Subject 71D | IGHV3-21 | IGHJ6 | CARREWMAAVESYDMDW | IGHG1 | F |
| 1 | Subject 71D | IGHV3-23 | IGHJ2 | CAKVCSSSTSCSNAFGYFDLW | IGHM | D |
| 1 | Subject 71D | IGHV3-23 | IGHJ3 | CAKGDYDYGSGYYSDAFDW | IGHG1 | D |
| 1 | Subject 71D | IGHV3-23 | IGHJ3 | CAKVTEYYYDSSGYGFGAFDIW | IGHM | F |
| 1 | Subject 71D | IGHV3-23 | IGHJ4 | CAKRDYDGGGYGIDYW | IGHM | C |
| 1 | Subject 71D | IGHV3-30 | IGHJ4 | CAKDFLTLYSSTWYGGPFDSW | IGHA1 | E |
| 1 | Subject 71D | IGHV3-30-3 | IGHJ4 | CARDYVDYFDYW | IGHG1 | F |
| 1 | Subject 71D | IGHV3-33 | IGHJ1 | CASAIAAAGTTFPQHW | IGHM | E |
| 1 | Subject 71D | IGHV3-33 | IGHJ4 | CARESYYYGPDYW | IGHM | C |
| 1 | Subject 71D | IGHV3-33 | IGHJ4 | CARDGSCSGSGSCYFGSLAYW | IGHM | C |
| 1 | Subject 71D | IGHV3-33 | IGHJ5 | CARESYYYGLDSG | ND | D |
| 1 | Subject 71D | IGHV3-33 | IGHJ5 | CARVGYSSSWWGWFDPW | IGHM | D |
| 1 | Subject 71D | IGHV3-7 | IGHJ4 | CARGGGNDYW | IGHM | E |
| 1 | Subject 71D | IGHV4-39 | IGHJ1 | CASRSASSWGWLHHW | IGHM | C |
| 1 | Subject 71D | IGHV4-39 | IGHJ4 | CARRGPWGGNWDFDYW | IGHM | E |
| 1 | Subject 71D | IGHV4-39 | IGHJ4 | CARYATTAVGGRSFDYW | IGHM | F |
| 1 | Subject 71D | IGHV4-39 | IGHJ4 | CARPAYVDAMVPYFDYW | IGHM | C |
| 1 | Subject 71D | IGHV4-39 | IGHJ5 | CARQGVRYNSGFYLNWFDPW | IGHG1 | E |
| 1 | Subject 71D | IGHV4-39 | IGHJ5 | CVRQGIHYDVVFRNLWFDSW | IGHG1 | F |
| 1 | Subject 71D | IGHV4-59 | IGHJ3 | CARDRTLYYSGGSANAFDIW | IGHM | F |
| 1 | Subject 71D | IGHV5-51 | IGHJ4 | CARHGPEGYFDYW | IGHM | D |
| 1 | Subject 71D | IGHV6-1 | IGHJ4 | CARGGAIFGWFDYW | IGHG1 | C |
| 1 | Subject 71D | IGHV7-4-1 | IGHJ6 | CARGYSSGWAYYYYMDW | IGHM | F |
| 1 | Subject 71E | IGHV1-18 | IGHJ4 | CARIYYDSSGYLFDYW | IGHM | D |
| 1 | Subject 71E | IGHV3-21 | IGHJ5 | CARDRRGYGGYGVGFDPW | IGHM | D |
| 1 | Subject 71E | IGHV3-23 | IGHJ3 | CVKDTNCPTECAFDFW | IGHM | C |
| 1 | Subject 71E | IGHV3-23 | IGHJ4 | CAKGPDSSTWSGSPPDYW | IGHM | C |
| 1 | Subject 71E | IGHV3-30 | IGHJ4 | CARLYDSSGEKSW | IGHM | D |
| 1 | Subject 71E | IGHV3-30 | IGHJ6 | CAREFRSDFWTGYSTNKAYYYGMDW | IGHG1 | C |
| 1 | Subject 71E | IGHV3-33 | IGHJ3 | CASLWGDWDADFIC | IGHM | F |
| 1 | Subject 71E | IGHV3-33 | IGHJ4 | CARDEGYW | IGHM | C |
| 1 | Subject 71E | IGHV3-33 | IGHJ4 | CAGGATTHYW | IGHM | C |
| 1 | Subject 71E | IGHV3-33 | IGHJ4 | CASKGGGFDYW | IGHM | C |
| 1 | Subject 71E | IGHV3-33 | IGHJ4 | CARDPATREGYW | IGHM | F |
| 1 | Subject 71E | IGHV3-33 | IGHJ6 | CARARDCSSTSCYPLDW | IGHM | D |
| 1 | Subject 71E | IGHV3-48 | IGHJ4 | CARAVGLWELLHFDW | IGHG1 | C |
| 1 | Subject 71E | IGHV3-7 | IGHJ4 | CSRESTGYIYGDCW | IGHM | D |
| 1 | Subject 71E | IGHV3-74 | IGHJ4 | CATTPGYFDYW | IGHM | C |
| 1 | Subject 71E | IGHV3-74 | IGHJ4 | CASRASPYGDPWDNW | IGHM | D |
| 1 | Subject 71E | IGHV4-34 | IGHJ5 | CARIGYSSSWYWFDPW | IGHM | F |
| 1 | Subject 71E | IGHV4-39 | IGHJ6 | CGRQVPYHGINW | IGHM | C |
| 1 | Subject 71E | IGHV4-61 | IGHJ4 | CARLVGGTFFDYW | IGHG1 | F |
| 1 | Subject 71E | IGHV5-51 | IGHJ4 | CARLEGSSSSCSYW | IGHM | F |

**Extended Data Table 3:** Summary of cloned CSP-specific BCRs gene usage and binding specificity as determined by ELISA.

| Ab ID | Subject | Original Isotype | Heavy Chain | Light Chain | IgG |  |  | IgM<br>CSP |
| --- | --- | --- | --- | --- | --- | --- | --- | --- |
|  |  |  |  |  | CSP | repeat | C-term |  |
| 82 | 71C | IgG | IGHV3-48 | IGKV3-1 | +++ | - | +++ | ND |
| 83 | 71C | IgM | IGHV3-33 | IGKV4-1 | - | - | - | +++ |
| 84 | 71C | IgM | IGHV2-5 | IGLV1-44 | - | - | - | - |
| 85 | 71C | IgM+B7 | IGHV3-21 | IGKV3-15 | +++ | - | +++ | ND |
| 86 | 71C | IgM | IGHV3-33 | IGKV3-1 | - | - | - | +++ |
| 87 | 71C | IgM | IGHV4-59 | IGLV3-1 | - | - | - | +++ |
| 88 | 71C | IgG | IGHV3-33 | IGKV4-1 | + | - | + | ND |
| 89 | 71C | IgG | IGHV3-48 | IGLV3-21 | +++ | - | +++ | ND |
| 90 | 71E | IgM | IGHV3-33 | IGKV1-5 | - | - | - | +++ |
| 91 | 71E | IgM | IGHV3-23 | IGLV1-39 | - | - | - | +++ |
| 92 | 71E | IgG | IGHV3-48 | IGLV3-21 | +++ | - | +++ | ND |
| 93 | 71C | IgG | IGHV3-30 | IGKV1-39 | +++ | - | +++ | ND |
| 94 | 71E | IgM | IGHV4-39 | IGKV3D-15 | - | - | - | ND |
| 95 | 71E | IgG | IGHV4-61 | IGKV1-33 | +++ | - | +++ | ND |
| 96 | 71E | IgM | IGHV4-34 | IGKV1-39 | - | - | - | - |
| 97 | 71E | IgM | IGHV5-51 | IGLV3-1 | - | - | - | +++ |
| 98 | 71D | IgM | IGHV1-3 | IGLV1-27 | +++ | + | - | +++ |
| 99 | 71D | IgG | IGHV1-24 | IGLV2-14 | - | - | - | ND |
| 100 | 71D | IgG | IGHV6-1 | IGKV1-8 | +++ | - | +++ | ND |
| 101 | 71D | IgA | IGHV3-23 | IGLV3-1 | +++ | - | +++ | + |
| 102 | 71D | IgG | IGHV4-39 | IGKV3-20 | +++ | - | +++ | ND |
| 103 | 71D | IgM | IGHV7-4-1 | IGKV2-28 | - | - | - | +++ |
| 104 | 71D | IgM | IGHV4-39 | IGKV1-5 | - | - | - | - |
| 105 | 71D | IgG | IGHV3-30-3 | IGKV3-11 | +++ | - | + | ND |
| 107 | 71C | IgA | IGHV3-48 | IGLV4-3 | +++ | + | - | +++ |
| 108 | 71C | IgA | IGHV3-33 | IGLV1-51 | - | - | - | +++ |
| 109 | 71D | IgA | IGHV1-3 | IGKV3-11 | - | - | - | +++ |
| 110 | 71D | IgA | IGHV3-30 | IGLV3-19 | - | - | - | - |
| 112 | 71C | IgG | IGHV3-33 | IGKV2-24 | - | - | - | ND |
| 113 | 71C | IgG | IGHV3-23 | IGLV3-1 | - | - | - | ND |
| 114 | 71E | IgG | IGHV3-30 | IGKV3-11 | +++ | - | +++ | ND |
| 115 | 71E | IgG | IGHV3-23 | IGLV3-1 | +++ | - | +++ | ND |
| 116 | 71D | IgG | IGHV3-30 | IGKV1-5 | +++ | +++ | - | ND |
| 118 | 71D | IgG | IGHV3-30 | IGKV1-5 | +++ | +++ | - | ND |

**Extended Data Table 4:** Summary of C-term specific antibodies tested in vitro and in vivo.

| Ab ID | Isotype | Heavy Chain | Light Chain | FL ELISA | Repeat | C-term | Competes 236 | Competes 1512 | Binds Spz | Selected for further testing | Kd (nM) |  | In vivo protection |
| --- | --- | --- | --- | --- | --- | --- | --- | --- | --- | --- | --- | --- | --- |
|  |  |  |  |  |  |  |  |  |  |  | CSP-FL | P16 |  |
| 82 | IgG | IGHV3-48 | IGKV3-1 | +++ | - | +++ | +++ | - | +++ |  |  |  |  |
| 85 | IgM | IGHV3-21 | IGKV3-15 | +++ | - | +++ | +++ | - | +++ | X | 14.6 | 19.8 | NO |
| 89 | IgG | IGHV3-48 | IGLV3-21 | +++ | - | +++ | +++ | - | +++ |  |  |  |  |
| 92 | IgG | IGHV3-48 | IGLV3-21 | +++ | - | +++ | +++ | - | +++ | X | 7.3 | 21.1 | NO |
| 93 | IgG | IGHV3-30 | IGKV1-39 | +++ | - | +++ | - | + | +++ | X | 52.4 | 11.4 | NO |
| 95 | IgG | IGHV4-61 | IGKV1-33 | +++ | - | +++ | - | + | +++ |  |  |  |  |
| 100 | IgG | IGHV6-1 | IGKV1-8 | +++ | - | +++ | - | - | +++ | X | 14.7 | 19.3 | NO |
| 101 | IgA | IGHV3-23 | IGLV3-1 | +++ | - | +++ | - | +++ | +++ | X | 3.46 | 1.3 | NO |
| 102 | IgG | IGHV4-39 | IGKV3-20 | +++ | - | +++ | - | - | +++ |  |  |  |  |
| 105 | IgG | IGHV3-30-3 | IGKV3-11 | +++ | - | + | - | - | +++ |  |  |  |  |
| 114 | IgG | IGHV3-30 | IGKV3-11 | +++ | - | +++ | ND | ND | +++ | X | 41.8 | 68.2 | NO |
| 115 | IgG | IGHV3-23 | IGLV3-1 | +++ | - | +++ | ND | ND | +++ |  |  |  |  |

**Extended Data Table 5:**
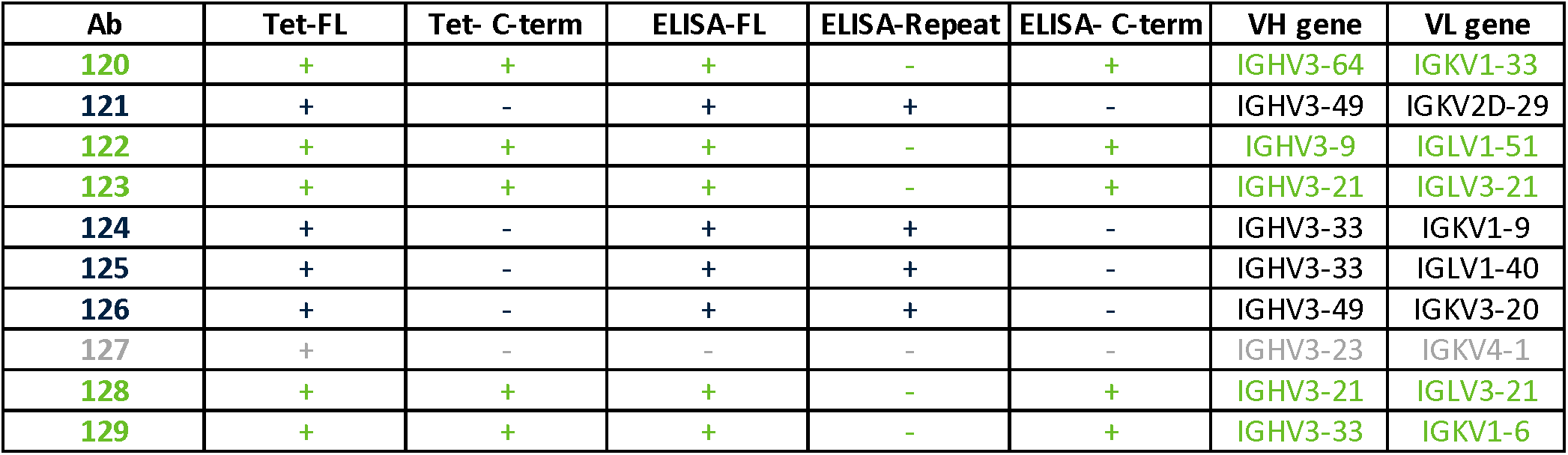
Comparison of tetramer and ELISA binding for dual tetramer approach.

| Ab | Tet-FL | Tet- C-term | ELISA-FL | ELISA-Repeat | ELISA- C-term | VH gene | VL gene |
| --- | --- | --- | --- | --- | --- | --- | --- |
| 120 | + | + | + | - | + | IGHV3-64 | IGKV1-33 |
| 121 | + | - | + | + | - | IGHV3-49 | IGKV2D-29 |
| 122 | + | + | + | - | + | IGHV3-9 | IGLV1-51 |
| 123 | + | + | + | - | + | IGHV3-21 | IGLV3-21 |
| 124 | + | - | + | + | - | IGHV3-33 | IGKV1-9 |
| 125 | + | - | + | + | - | IGHV3-33 | IGLV1-40 |
| 126 | + | - | + | + | - | IGHV3-49 | IGKV3-20 |
| 127 | + | - | - | - | - | IGHV3-23 | IGKV4-1 |
| 128 | + | + | + | - | + | IGHV3-21 | IGLV3-21 |
| 129 | + | + | + | - | + | IGHV3-33 | IGKV1-6 |

## References

1. World malaria report 2025. https://www.who.int/teams/global-malaria-programme/reports/world-malaria-report-2025.

2. Kayentao, K. et al. Safety and Efficacy of a Monoclonal Antibody against Malaria in Mali. New England Journal of Medicine 387, 1833–1842 (2022).

3. Lyke, K. E. et al. Low-dose intravenous and subcutaneous CIS43LS monoclonal antibody for protection against malaria (VRC 612 Part C): a phase 1, adaptive trial. Lancet Infect. Dis. 23, 578– 588 (2023).

4. Wu, R. L. et al. Low-Dose Subcutaneous or Intravenous Monoclonal Antibody to Prevent Malaria. New England Journal of Medicine 387, 397–407 (2022).

5. Olotu, A. et al. Seven-Year Efficacy of RTS,S/AS01 Malaria Vaccine among Young African Children. N. Engl. J. Med. 374, 2519–2529 (2016).

6. Olotu, A. et al. Efficacy of RTS,S/AS01E malaria vaccine and exploratory analysis on anti-circumsporozoite antibody titres and protection in children aged 5-17 months in Kenya and Tanzania: a randomised controlled trial. Lancet Infect. Dis. 11, 102–109 (2011).

7. Agnandji, S. T. et al. Efficacy and Safety of the RTS,S/AS01 Malaria Vaccine during 18 Months after Vaccination: A Phase 3 Randomized, Controlled Trial in Children and Young Infants at 11 African Sites. PLoS Med. 11, e1001685 (2014).

8. Clinical Trials Partnership, S. Efficacy and safety of RTS,S/AS01 malaria vaccine with or without a booster dose in infants and children in Africa: Final results of a phase 3, individually randomised, controlled trial. The Lancet 386, 31–45 (2015).

9. White, M. T. et al. Immunogenicity of the RTS,S/AS01 malaria vaccine and implications for duration of vaccine efficacy: secondary analysis of data from a phase 3 randomised controlled trial. Lancet Infect. Dis. 15, 1450–1458 (2015).

10. Stoute, J. A. et al. A preliminary evaluation of a recombinant circumsporozoite protein vaccine against Plasmodium falciparum malaria. RTS,S Malaria Vaccine Evaluation Group. N. Engl. J. Med. 336, 86–91 (1997).

11. Clinical Trials Partnership, S. Efficacy and safety of RTS,S/AS01 malaria vaccine with or without a booster dose in infants and children in Africa: Final results of a phase 3, individually randomised, controlled trial. The Lancet 386, 31–45 (2015).

12. Kurtovic, L., Drew, D. R., Dent, A. E., Kazura, J. W. & Beeson, J. G. Antibody Targets and Properties for Complement-Fixation Against the Circumsporozoite Protein in Malaria Immunity. Front. Immunol. 12, 775659 (2021).

13. Sanchez, L. et al. Antibody responses to the RTS, S/AS01E vaccine and Plasmodium falciparum antigens after a booster dose within the phase 3 trial in Mozambique. npj Vaccines 2020 *5:1* 5, 46- (2020).

14. Ubillos, I. et al. Baseline exposure, antibody subclass, and hepatitis B response differentially affect malaria protective immunity following RTS,S/AS01E vaccination in African children. BMC Medicine 2018 *16:1* **16**, 197- (2018).

15. Dobaño, C. et al. Concentration and avidity of antibodies to different circumsporozoite epitopes correlate with RTS,S/AS01E malaria vaccine efficacy. Nature Communications 2019 *10:1* **10**, 1–13 (2019).

16. Lara-Escandell, M. et al. Duration and association with protection of NANP-repeat-specific and C-terminus-specific anti-circumsporozoite protein IgG responses following RTS,S/AS01E vaccination: an observational ancillary immunological study of a phase 3 clinical trial. Lancet Infect. Dis. https://doi.org/10.1016/S1473-3099(26)00081-2 (2026) doi:10.1016/S1473-3099(26)00081-2.

17. Cockburn, I. A. & Seder, R. A. Malaria prevention: from immunological concepts to effective vaccines and protective antibodies. Nat. Immunol. 19, 1199–1211 (2018).

18. Schofield, L. On the function of repetitive domains in protein antigens of Plasmodium and other eukaryotic parasites. Parasitol. Today 7, 99–105 (1991).

19. Schofield, L. & Uadia, P. Lack of Ir gene control in the immune response to malaria. I. A thymus-independent antibody response to the repetitive surface protein of sporozoites. The Journal of Immunology 144, 2781–2788 (1990).

20. Chatterjee, D. et al. Avid binding by B cells to the Plasmodium circumsporozoite protein repeat suppresses responses to protective subdominant epitopes. Cell Rep. 35, 108996 (2021).

21. Oludada, O. E. et al. Molecular and functional properties of human Plasmodium falciparum CSP C-terminus antibodies . EMBO Mol. Med. 15, (2023).

22. Scally, S. W. et al. Rare PfCSP C-terminal antibodies induced by live sporozoite vaccination are ineffective against malaria infection. Journal of Experimental Medicine 215, 63–75 (2018).

23. Beutler, N. et al. A novel CSP C-terminal epitope targeted by an antibody with protective activity against Plasmodium falciparum. PLoS Pathog. 18, (2022).

24. Regules, J. A. et al. Fractional Third and Fourth Dose of RTS,S/AS01 Malaria Candidate Vaccine: A Phase 2a Controlled Human Malaria Parasite Infection and Immunogenicity Study. J. Infect. Dis. 214, 762–771 (2016).

25. Moon, J. E. et al. A Phase IIa Controlled Human Malaria Infection and Immunogenicity Study of RTS,S/AS01E and RTS,S/AS01_B_ Delayed Fractional Dose Regimens in Malaria-Naive Adults. J. Infect. Dis. 222, 1681–1691 (2020).

26. Krishnamurty, A. T. et al. Somatically Hypermutated Plasmodium-Specific IgM(+) Memory B Cells Are Rapid, Plastic, Early Responders upon Malaria Rechallenge. Immunity 45, 402–414 (2016).

27. Oyen, D. et al. Structural basis for antibody recognition of the NANP repeats in Plasmodium falciparum circumsporozoite protein. Proc. Natl. Acad. Sci. U. S. A. 114, E10438–E10445 (2017).

28. Kisalu, N. K. et al. A human monoclonal antibody prevents malaria infection by targeting a new site of vulnerability on the parasite. Nat. Med. 24, 408–416 (2018).

29. Imkeller, K. et al. Antihomotypic affinity maturation improves human B cell responses against a repetitive epitope. Science 360, 1358–1362 (2018).

30. Triller, G. et al. Natural Parasite Exposure Induces Protective Human Anti-Malarial Antibodies. Immunity 47, 1197–1209.e10 (2017).

31. Zavala, F., Cochrane, A. H., Nardin, E. H., Nussenzweig, R. S. & Nussenzweig, V. Circumsporozoite proteins of malaria parasites contain a single immunodominant region with two or more identical epitopes. J. Exp. Med. 157, 1947–1957 (1983).

32. Beeson, J. G. et al. Challenges and strategies for developing efficacious and long-lasting malaria vaccines. Sci. Transl. Med. 11, (2019).

33. Pholcharee, T. et al. Structural and biophysical correlation of anti-NANP antibodies with in vivo protection against P. falciparum. Nature Communications 2021 12:1 *12*, 1–14 (2021).

34. Oyen, D. et al. Structural basis for antibody recognition of the NANP repeats in Plasmodium falciparum circumsporozoite protein. Proc. Natl. Acad. Sci. U. S. A. 114, E10438–E10445 (2017).

35. Murugan, R. et al. Evolution of protective human antibodies against Plasmodium falciparum circumsporozoite protein repeat motifs. Nature Medicine 2020 26:*7* *26*, 1135–1145 (2020).

36. Flores-Garcia, Y. et al. Optimization of an in vivo model to study immunity to Plasmodium falciparum pre-erythrocytic stages. Malar. J. 18, (2019).

37. Raghunandan, R. et al. Characterization of two in vivo challenge models to measure functional activity of monoclonal antibodies to Plasmodium falciparum circumsporozoite protein. Malar. J. 19, 1–15 (2020).

38. Seaton, K. E. et al. Subclass and avidity of circumsporozoite protein specific antibodies associate with protection status against malaria infection. npj Vaccines 2021 *6:1* **6**, 1–13 (2021).

39. Young, W. C. et al. Comprehensive Data Integration Approach to Assess Immune Responses and Correlates of RTS,S/AS01-Mediated Protection From Malaria Infection in Controlled Human Malaria Infection Trials. Front. Big Data 4, (2021).

40. Das, J. et al. Delayed fractional dosing with RTS,S/AS01 improves humoral immunity to malaria via a balance of polyfunctional NANP6- and Pf16-specific antibodies. Med 2, 1269-1286.e9 (2021).

41. Suscovich, T. J. et al. Mapping functional humoral correlates of protection against malaria challenge following RTS,S/AS01 vaccination. Sci. Transl. Med. 12, 4757 (2020).

42. Wang, L. T. et al. A Potent Anti-Malarial Human Monoclonal Antibody Targets Circumsporozoite Protein Minor Repeats and Neutralizes Sporozoites in the Liver. Immunity 53, 733–744.e8 (2020).

43. Gaudinski, M. R. et al. A Monoclonal Antibody for Malaria Prevention. N. Engl. J. Med. 385, 803– 814 (2021).

44. Williams, K. L. et al. A candidate antibody drug for prevention of malaria. Nature Medicine 2024 30:*1* 30, 117–129 (2024).

45. Ali, M. S. et al. The anti-circumsporozoite antibody response to repeated, seasonal booster doses of the malaria vaccine RTS,S/AS01E. npj Vaccines 2025 10*:1* **10**, 1–11 (2025).

46. McNamara, H. A. et al. Antibody Feedback Limits the Expansion of B Cell Responses to Malaria Vaccination but Drives Diversification of the Humoral Response. Cell Host Microbe 28, 572–585.e7 (2020).

47. Chaudhury, S. et al. Breadth of humoral immune responses to the C-terminus of the circumsporozoite protein is associated with protective efficacy induced by the RTS,S malaria vaccine. Vaccine 39, 968–975 (2021).

48. Feldmann, M. & Basten, A. The relationship between antigenic structure and the requirement for thymus-derived cells in the immune response. J. Exp. Med. 134, 103–119 (1971).

49. Kato, Y. et al. Multifaceted Effects of Antigen Valency on B Cell Response Composition and Differentiation In Vivo. Immunity 53, 548–563.e8 (2020).

50. O’connor, B. P. et al. Imprinting the Fate of Antigen-Reactive B Cells through the Affinity of the B Cell Receptor. J Immunol 177, 7723 (2006).

51. Ochiai, K. et al. Transcriptional regulation of germinal center B and plasma cell fates by dynamical control of IRF4. Immunity 38, 918–929 (2013).

52. Paus, D. et al. Antigen recognition strength regulates the choice between extrafollicular plasma cell and germinal center B cell differentiation. J. Exp. Med. 203, 1081–1091 (2006).

53. Schwickert, T. A. et al. A dynamic T cell-limited checkpoint regulates affinity-dependent B cell entry into the germinal center. Journal of Experimental Medicine 208, 1243–1252 (2011).

54. Papadatou, I., Tzovara, I. & Licciardi, P. V. The Role of Serotype-Specific Immunological Memory in Pneumococcal Vaccination: Current Knowledge and Future Prospects. Vaccines (Basel). 7, (2019).

55. Clutterbuck, E. A. et al. Pneumococcal Conjugate and Plain Polysaccharide Vaccines Have Divergent Effects on Antigen-Specific B Cells. J. Infect. Dis. 205, 1408 (2012).

56. Fisher, C. R. et al. T-dependent B cell responses to Plasmodium induce antibodies that form a high-avidity multivalent complex with the circumsporozoite protein. PLoS Pathog. 13, e1006469 (2017).

57. Ogieuhi, I. J. et al. A narrative review of the RTS S AS01 malaria vaccine and its implementation in Africa to reduce the global malaria burden. Discover Public Health 2024 21:*1* 21, 1–13 (2024).

58. Rodda, L. B. et al. Functional SARS-CoV-2-Specific Immune Memory Persists after Mild COVID-19. Cell 184, 169–183.e17 (2021).

59. Thouvenel, C. D. et al. Multimeric antibodies from antigen-specific human IgM+ memory B cells restrict Plasmodium parasites. J. Exp. Med. 218, (2021).

60. Hale, M. et al. IgM antibodies derived from memory B cells are potent cross-variant neutralizers of SARS-CoV-2. J. Exp. Med. 219, (2022).

61. Hale, M. et al. Monoclonal antibodies derived from B cells in subjects with cystic fibrosis reduce Pseudomonas aeruginosa burden in mice. Elife 13, (2024).

62. Zheng, Z. et al. Anchored multiplex PCR for targeted next-generation sequencing. Nature Medicine 2014 20:*12* 20, 1479–1484 (2014).

63. Alamyar, E., Duroux, P., Lefranc, M. P. & Giudicelli, V. IMGT(®) tools for the nucleotide analysis of immunoglobulin (IG) and T cell receptor (TR) V-(D)-J repertoires, polymorphisms, and IG mutations: IMGT/V-QUEST and IMGT/HighV-QUEST for NGS. Methods Mol. Biol. 882, 569–604 (2012).

64. Tiller, T. et al. Efficient generation of monoclonal antibodies from single human B cells by single cell RT-PCR and expression vector cloning. J. Immunol. Methods 329, 112–124 (2008).

